# Mechanistic origin of drug interactions between translation-inhibiting antibiotics

**DOI:** 10.1101/843920

**Authors:** Bor Kavčič, Gašper Tkačik, Tobias Bollenbach

**Affiliations:** IST Austria, 3400 Klosterneuburg, Austria; Institute for Biological Physics, University of Cologne, 50937 Cologne, Germany

**Keywords:** antibiotics, drug combinations, drug interaction mechanisms, growth laws, bacterial physiology, translation, translation inhibitors, ribosomes, translation factors, ribosome traffic jams

## Abstract

Antibiotics that interfere with translation, when combined, interact in diverse and difficult-to-predict ways. Here, we demonstrate that these interactions can be accounted for by “translation bottlenecks”: points in the translation cycle where antibiotics block ribosomal progression. To elucidate the underlying mechanisms of drug interactions between translation inhibitors, we generated translation bottlenecks genetically using inducible control of translation factors that regulate well-defined translation cycle steps. These perturbations accurately mimicked antibiotic action and their interactions, supporting that the interplay of different translation bottlenecks causes these interactions. We further showed that the kinetics of drug uptake and binding together with growth laws allows direct prediction of a large fraction of observed interactions, yet fails for suppression. Simultaneously varying two translation bottlenecks in the same cell revealed how the dense traffic of ribosomes and competition for translation factors results in previously unexplained suppression. This result highlights the importance of “continuous epistasis” in bacterial physiology.

## 1 Introduction

Inhibiting translation is one of the most common antibiotic modes of action, crucial for restraining pathogenic bacteria [Walsh, 2003]. Antibiotics targeting translation interfere with either the assembly or the processing of the ribosome, or with the proper utilization of charged tRNAs and translation factors (Fig. 1A,B; Table 1) [Wilson, 2014]. Still, the exact modes of action and physiological responses to many such translation inhibitors are less clear, and responses to drug combinations are even harder to understand, even though they offer effective ways of fighting antibiotic resistance [Yeh *et al.*, 2009]. Recently, mechanism-independent mathematical approaches to predict the responses to multi-drug combinations were proposed [Zimmer *et al.*, 2016; Wood *et al.*, 2012], yet these approaches rely on prior knowledge of pairwise drug interactions, which are diverse and have notoriously resisted prediction. They include synergism (inhibition is stronger than predicted), antagonism (inhibition is weaker), and suppression (one of the drugs loses potency) [Bollenbach, 2015; Mitosch and Bollenbach, 2014] (Fig. 1C). To design optimized treatments, the ability to predict or alter drug interactions is crucial – a challenge that would be facilitated by understanding their underlying mechanisms [Chevereau and Bollenbach, 2015].

**Table 1:**
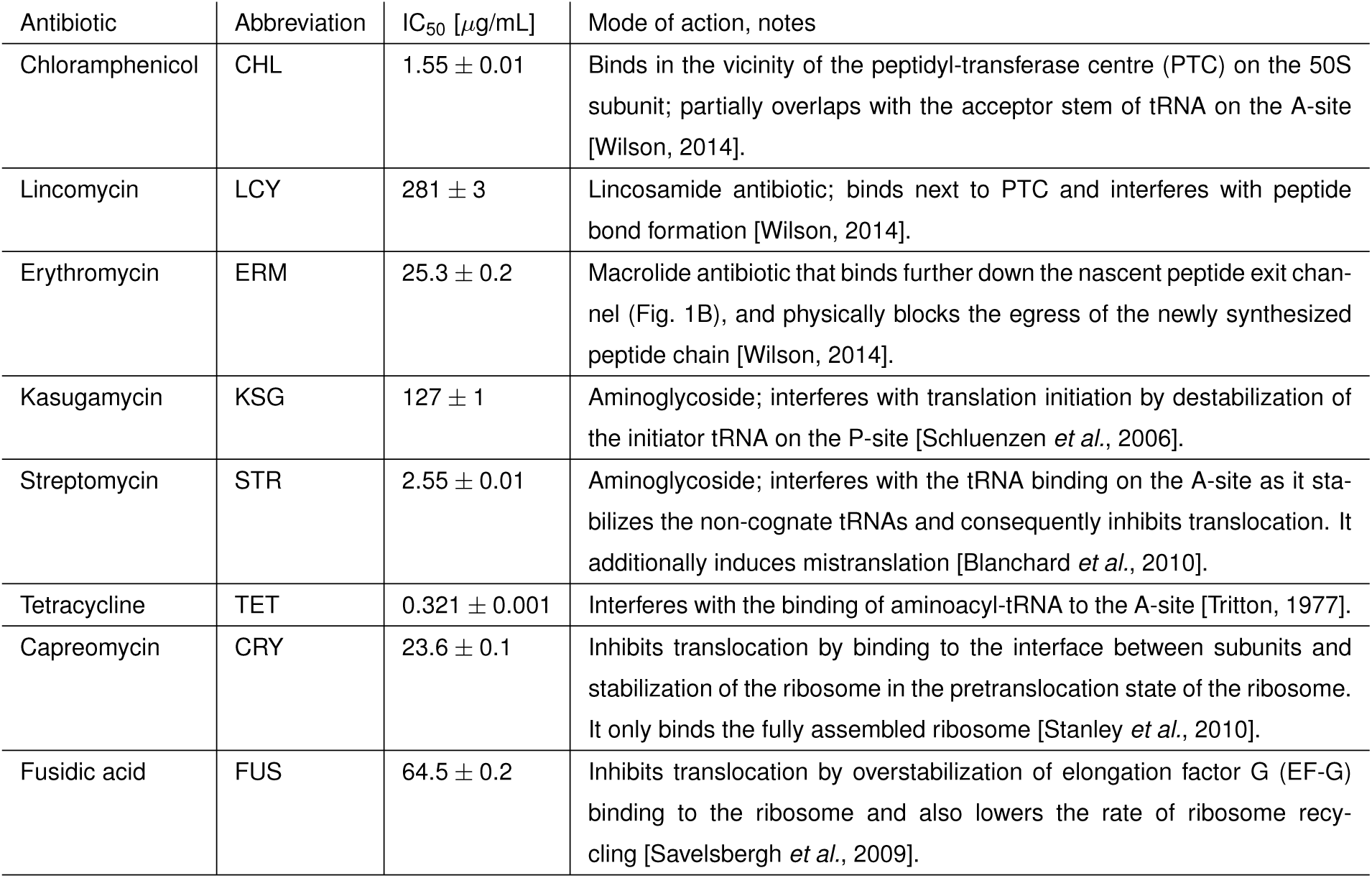
Translation-targeting antibiotics used in this study and their characteristics.

**Figure 1:**
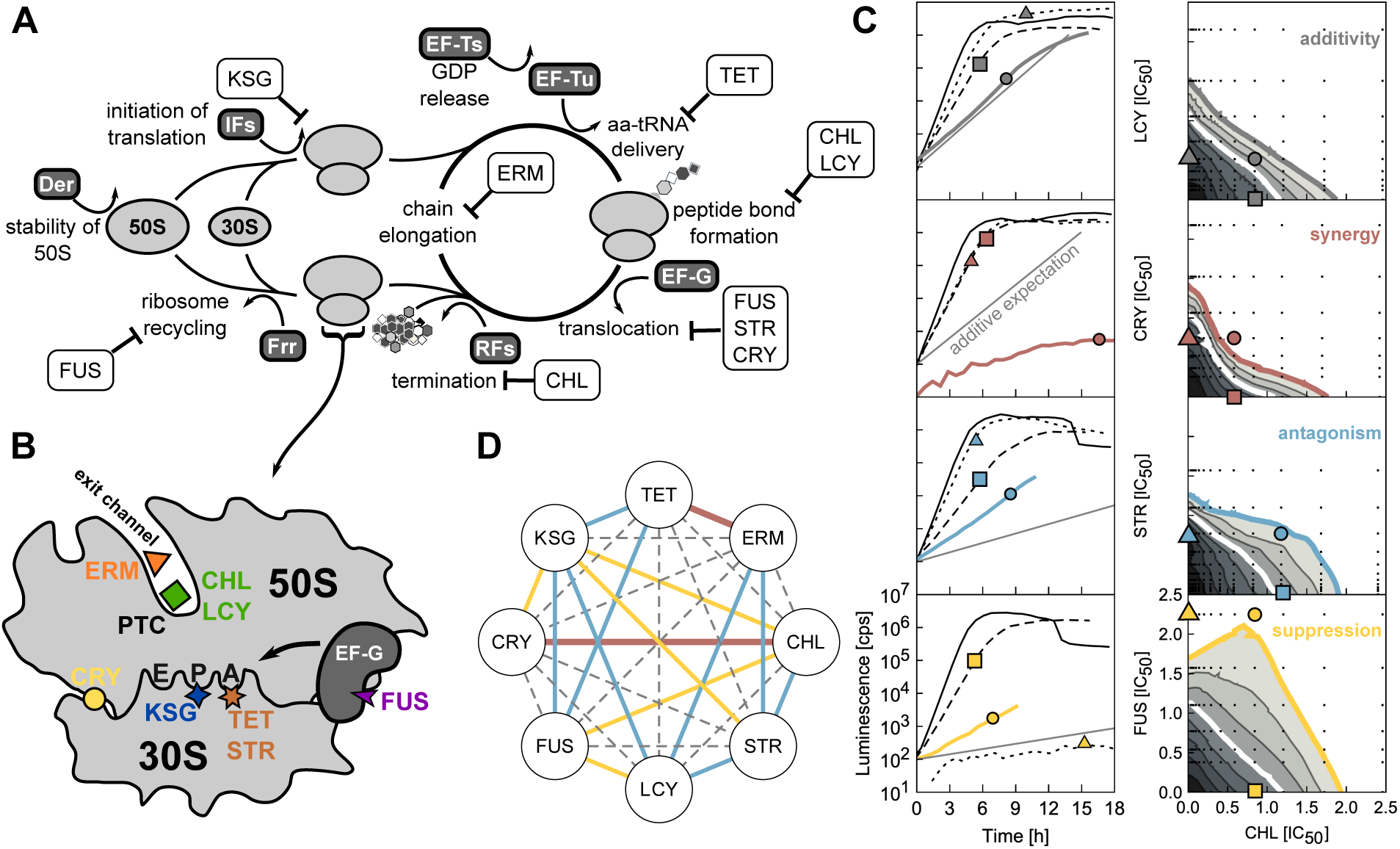
Antibiotics targeting different translation steps show diverse drug interactions. **(A,B)** Schematic of the translation cycle and translation inhibitors. Translation factors are shown in dark gray boxes. Stability of the large subunit is mediated by Der and initiation by initiation factors (IFs). Elongation factors Tu and G (EF-Tu, EF-G) catalyze ribosome progression. Release of GDP from EF-Tu is facilitated by EF-Ts. Release factors (RFs) facilitate the ejection of the finished peptide from the ribosome, whose recycling is mediated by the factor for ribosome recycling (Frr). Translation inhibitors are shown in white boxes (abbreviations in Table 1). **(C)** Examples of growth curves obtained by luminescence assay (left column) in the presence of different antibiotics and their combinations and response surfaces corresponding to different interaction types (right column) (Methods). Symbols on the growth curves indicate the condition used: no symbol, triangle, square and a circle correspond to no drug, CHL-only, second drug only (see vertical axis), and the combination of both, respectively. The growth curves were shifted in time so as to originate from the same point at time 0. Drug interactions are determined based on the shape of lines of equal growth (isoboles). If the addition of the second drug has the same effect as increasing the concentration of the first, the isoboles are straight lines [Loewe and Muischnek, 1926]. Deviations from this additive expectation reveal synergism (the combined effect is stronger and isoboles curve towards the origin), antagonism (the effect is weaker and isoboles curve away from the origin), or suppression (at least one of the drugs loses potency due to the other). **(D)** The drug-interaction network of translation inhibitors. Color-code is as in (C); dashed gray lines denote additivity.

Apart from their clinical relevance, antibiotic combinations provide powerful, quantitative and controlled means of studying perturbations of cell physiology [Falconer *et al.*, 2011] – conceptually similar to studies of epistasis between double gene knockouts [Yeh *et al.*, 2006; Segre *et al.*, 2005]. Translation inhibitors are particularly suited for this purpose since translation is a fundamental, yet complex multi-step process that still lacks a comprehensive quantitative description. Part of any such description are “growth laws,” which quantitatively capture the compensatory upregulation of the translational machinery in response to perturbations of translation [Scott *et al.*, 2010]. Growth laws have enabled a model that elegantly explains the growth-dependent bacterial susceptibility to individual translation inhibitors [Greulich *et al.*, 2015]. Finally, well defined translation steps cannot only be perturbed chemically [Blanchard *et al.*, 2010; Wilson, 2014], but also genetically, as these steps are regulated by translation factors – specialized proteins that mediate the stability of ribosomal subunits, catalyze assembly of 70S ribosome and initiation, deliver charged tRNAs to the ribosome, release finished peptides, and mediate ribosome recycling (Fig. 1A). Both genetic and chemical perturbations obstruct the progression of ribosomes along the translation cycle, which generally results in a lower growth rate. Comparing the effects of antibiotics to those of precisely defined genetic perturbations offers an opportunity to elucidate the mechanisms responsible for drug interactions between translation inhibitors.

As drug interactions are largely determined by the modes of action of the combined antibiotics [Yeh *et al.*, 2006], we hypothesized that a key determinant of interactions between pairs of translation inhibitors are the specific steps in the translation cycle where the two inhibitors halt ribosomal progression (Fig. 1A). As a second key determinant of these drug interactions, we considered the compensatory physiological response to translation inhibition captured quantitatively by ribosomal growth laws [Scott *et al.*, 2010] together with the kinetics of antibiotic transport and ribosome binding. We show that these determinants suffice to understand how most drug interactions between translation inhibitors emerge and that they can be predicted solely from known responses to the individual drugs. To establish this result, we used a combination of precise growth measurements, quantitative genetic perturbations of the translation machinery, and theoretical modeling.

## 2 Results

### 2.1 Pairwise interactions between translation inhibitors are highly diverse

To systematically map the network of drug interactions between translation inhibitors, we selected eight representative antibiotics that interfere with different stages of translation and bind to different sites on the ribosome (Fig. 1A,B; Table 1). To this end, we determined high-resolution dose-response surfaces for all pairwise combinations of these antibiotics (Fig. 1C), by measuring growth rates in two-dimensional drug concentration matrices using a highly precise technique based on bioluminescence [Kishony and Leibler, 2003; Yeh *et al.*, 2006; Chait *et al.*, 2007] (Methods). To quantify the drug interaction, we defined the Loewe interaction score *LI* that integrates deviations from Loewe additivity (Fig. 1C, Methods). In this way, we characterized all twenty-eight pairwise interactions and constructed the interaction network between the translation inhibitors (Fig. 1D).

The translation inhibitor interaction network (Fig. 1D) that we measured has several notable properties. First, antibiotics with similar mode of action tend to exhibit additive drug interactions: in particular, there are purely additive interactions between capreomycin (CRY), fusidic acid (FUS), and streptomycin (STR) (which all inhibit translocation) and chloramphenicol (CHL) and lincomycin (LCY) (which both inhibit peptide bond formation), respectively. This observation is consistent with the view that drugs with similar mode of action can substitute for one another. Second, kasugamycin (KSG) is a prominent hub in the network: it shows almost exclusively antagonistic and suppressive interactions with other translation inhibitors. Third, we identified a previously unreported synergy between CRY and CHL. Some of the observed general trends in the drug interaction network, in particular the prevalence of antagonism, may be explained by a general physiological response to translation inhibition.

A number of the interactions we measured confirm previous reports. For example, synergy between erythromycin (ERM) and tetracycline (TET) was observed before [Yeh *et al.*, 2006; Russ and Kishony, 2018]. Additivity between CHL and TET was also reported; moreover, this interaction proved to be highly robust to genetic perturbations [Chevereau and Bollenbach, 2015]. Globally, antagonism and suppression are more common in the translation inhibitor interaction network than synergy, consistent with a general prevalence of antagonistic interactions between antibiotics [Brochado *et al.*, 2018].

### 2.2 Growth-law based biophysical model correctly predicts some interactions but fails to predict suppression

As a first step toward understanding the origin of the observed drug interactions, we developed a mathematical model that predicts such interactions from the effects of the individual drugs alone. We generalized a biophysical model for the effect of a single antibiotic on bacterial growth [Greulich *et al.*, 2015] to the situation where two antibiotics are present simultaneously. The model consists of ordinary differential equations taking into account passive antibiotic transport into the cell, binding to the ribosome (Fig. 2A,B), dilution of all molecular species due to cell growth, and the physiological response of the cell to the perturbation (Fig. 2C). The latter is described by ribosomal growth laws [Scott *et al.*, 2010; Greulich *et al.*, 2015], which quantitatively connect the growth rate to the total abundance of ribosomes when growth rate is varied by the nutrient quality of media or by translation inhibitors. All parameters of the model can be inferred from the dose-response curves of individual drugs (Fig. 2D).

**Figure 2:**
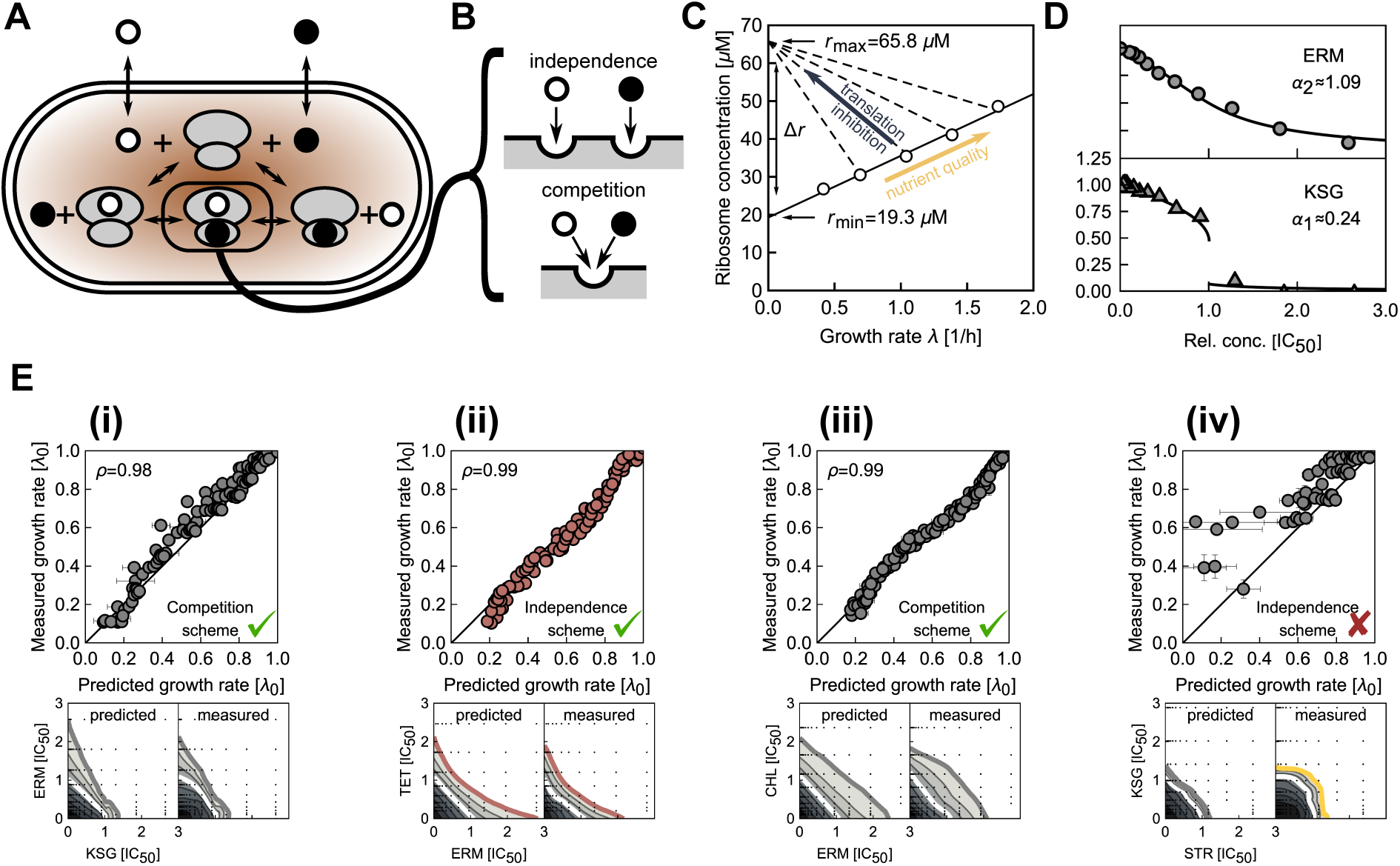
Mathematical model of combined antibiotic action based on growth laws partially predicts drug interactions. **(A)** Schematic of antibiotic binding and transport into the cell. Antibiotics (circles) bind to the unbound ribosomes (gray) in the first binding step; bound ribosomes can be bound by a second antibiotic. **(B)** Schematic of antibiotics binding independently (top) or competing for the same binding site (bottom). **(C)** Growth laws link intracellular ribosome concentration to the growth rate. Solid line: ribosome concentration when growth rate is varied by varying nutrient quality; dashed lines: ribosome concentration when growth rate is lowered by perturbation of translation. Circles show data from Ref. [Greulich *et al.*, 2015]. **(D)** Data points are dose-response curves for ERM and KSG; lines show best fits of the mathematical model. The best-fit values of the steepness parameter *α* that encapsulates kinetic and physiological parameters (Methods) are shown. Both shallow (top panel, ERM) and steep (bottom panel, KSG) dose-response curves are observed. **(E)** Examples of predicted dose-response surfaces. Scatter plot depicts correlation between predicted and measured growth rate. The binding scheme assumed is indicated on the bottom right and Pearson’s *ρ* on the top left. Predicted and measured dose-response surfaceare shown below the scatter plot. Color of 20% isobole (bottom) and plot markers (top) denotes the type of predicted interaction. **(i)** Response surface for antibiotics from (D). Here, the independent binding scheme quantitatively predicts the response surface. **(ii)** As in (i) but for ERM-TET; model with independent biding scheme correctly predicts mild synergism. **(iii)** For CHL-ERM, a competitive biding scheme results in an additive interaction, which is observed experimentally. **(iv)** Interaction between STR-KSG is not explained by the model.

When two different antibiotics are present simultaneously, separate variables are needed to describe ribosomes that are bound by either of the antibiotics individually or simultaneously by both (Fig. 2A). In the absence of knowledge about direct molecular interactions on the ribosome (as for the pairs of lankamycin and lankacidin or of dalfopristin and quinupristin [Harms *et al.*, 2004; Belousoff *et al.*, 2011]), we assumed that the antibiotic binding and unbinding rates are independent of any previously bound antibiotic (Fig. 2B). The resulting model makes direct predictions for drug interactions between translation inhibitors using only parameters that are inferred from the individual drug dose-response curves.

Using this model, we calculated the predicted response surfaces for all translation inhibitor pairs and compared them to the experimentally measured surfaces (Methods, Fig. 2E). Certain drug interactions were correctly predicted by this approach (ERM-KSG, TET-ERM in Fig. 2E-i and ii), indicating that binding kinetics and growth physiology alone suffice to explain these interactions. Correctly predicted drug interactions include additive cases which often involve antibiotics that have either the same mode of action (CRY-FUS, CRY-STR, FUS-STR, CHL-LCY) or partially overlapping binding sites (CHL-LCY, ERM-CHL) [Wilson, 2014]. For the latter, the assumption that the formation of the doubly-bound ribosome population is prohibited, which yields an additive response surface, offers even better agreement with the experimental data (Fig. 2E-iii).

Other drug interactions clearly deviated from the model predictions. An example is the suppressive/antagonistic interaction between STR and KSG, which was predicted to be additive (Fig. 2E-iv). Such clear deviations could originate from the direct molecular interactions of the drugs on the ribosome, and thus be specific for every pair of drugs. Alternatively, these mechanisms could originate from the multi-step structure of the translation cycle itself, making general predictions possible. In the most complex cases, drug interactions could result from drug effects that are unrelated to the primary drug target [Chevereau and Bollenbach, 2015], in particular from effects on drug uptake or efflux [Lazar *et al.*, 2013]. We focused on the most general hypothesis that drug interactions arise from the interplay of ribosomes halted in different stages of translation cycle such as initiation, translocation, recycling, etc. (Fig. 1).

### 2.3 Inducible genetic bottlenecks in translation strongly affect antibiotic efficacy

To test this hypothesis, we developed a technique to determine how halting ribosomes in different stages of the translation cycle affects the efficacy of various antibiotics. Specifically, we imposed artificial bottlenecks in translation by genetically limiting the expression of translation factors that catalyze well-defined translation steps [Cole *et al.*, 1987]. We constructed *E. coli* strains with translation factor genes under inducible control of a synthetic promoter [Lutz and Bujard, 1997]. These genes were integrated in the chromosome outside of their endogenous loci and the endogenous copy of the gene was disrupted (Fig. 3A; Methods). This yielded six strains that enable continuous control of key translation processes (Fig. 3B): stabilization of the 50S subunit (*der*), initiation (*infB*), delivery of charged tRNAs (*tufA/B*), release of GDP from elongation factors (*tsf*), translocation (*fusA*) and recycling of the ribosomes (*frr*) [Rodnina, 2018]. Reducing translation factor expression by varying the inducer concentration resulted in a gradual decrease in growth which stopped at almost complete cessation of growth, reflecting the essentiality of translation factors (Fig. 3C, Methods and SI). Since the endogenous regulation of translation factors generally follows that of the translation machinery [Maaløe, 1979; Gordon, 1970; Blumenthal *et al.*, 1976; Furano and Wittel, 1975], limiting the expression of a single translation factor imposes a highly specific bottleneck as all other components get upregulated. Furthermore, any global feedback regulation is left intact as we removed the factor from its native operon. These synthetic strains thus offer precise control over artificial translation bottlenecks that determine the rates of different translation steps.

**Figure 3:**
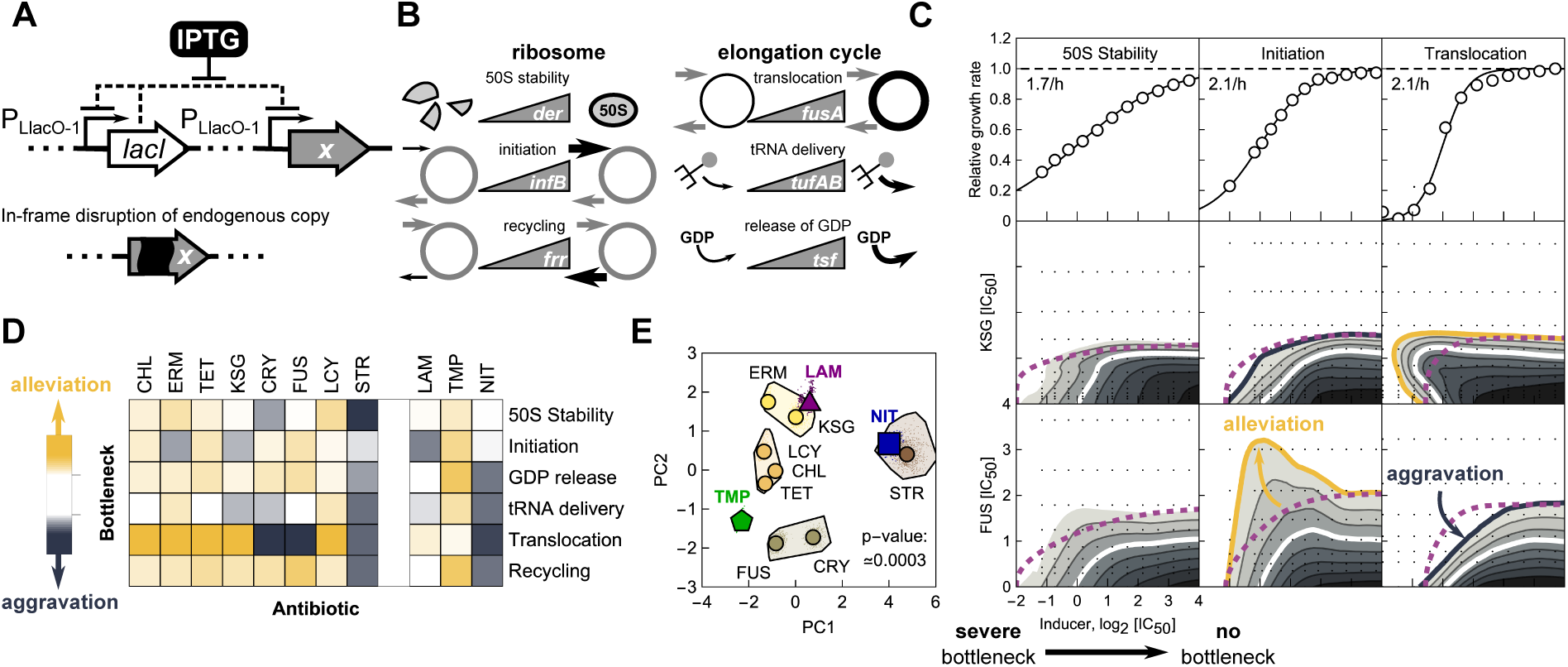
Artificial translation bottlenecks strongly affect antibiotic efficacy. **(A)** Schematic of synthetic regulation introduced to control the expression of a translation factor x, which creates an artificial bottleneck in translation at a well-defined stage; *lacI* codes for the Lac repressor, which represses the P_LlacO-1_-promoter (Methods, [Lutz and Bujard, 1997]). **(B)** Constructs were made for six translation factors mediating 50S stability (*der*), initiation (*infB*), recycling (*frr*), translocation (*fusA*), tRNA delivery (*tufAB*) and GDP release (*tsf*), respectively. Higher expression alleviates the artificial bottleneck. Thicker lines or arrows indicate higher rates. **(C)** Translation factor induction curves (upper row) and response surfaces over inducer-antibiotic grid for different antibiotics (KSG and FUS, middle and bottom row, respectively) in combination with different bottlenecks (50S stability, initiation, and translocation). Full induction of the translation factor rescues wild type growth; increasing bottleneck severity leads to smooth decrease in growth rate to zero. Comparison of the response surfaces with independent expectation (dashed purple line) identify alleviation (orange line) or aggravation (blue line). **(D)** Columns show bottleneck dependency vectors in color code; dependency vectors quantify the response of a given antibiotic to the translation bottlenecks (Methods). **(E)** Clustering of the bottleneck dependency vectors upon dimensionality-reduction by Principal Component Analysis (PCA; Methods). Circles show dependency vectors projected onto the first two principal components (PC1, PC2); colors indicate cluster identity. The extended cluster areas shown are convex hulls of bootstrapped projections (denoted by dots; Methods). Deviations of the three additional antibiotics LAM, NIT, and TMP are denoted by a purple triangle, blue square, and green pentagon, respectively. The observed clustering is highly significant (*p* ≈ 3 × 10^−4^, bootstrap; Methods).

We next used these strains to assess the impact of bottlenecks on antibiotic efficacy. Accordingly, we measured growth rates over a two-dimensional matrix of concentrations of inducer and antibiotic for each of the six strains (Fig. 3C; Methods). To address if the action of the antibiotic is independent of the translation bottleneck, we analyzed these experiments using a multiplicative null expectation. Note that additivity as used for antibiotics (Fig. 1C) is not a suitable null expectation here since the responses to increasing concentrations of antibiotic and inducer are opposite. However, if antibiotic action is independent of the translation bottleneck, the growth rate should be a product of the relative growth rates of each of the two perturbations acting individually. Independence implies that the dose-response surface is obtained as a multiplication of the antibiotic dose-response and the translation factor induction curve. Deviations from independence indicate a nontrivial interaction between the bottleneck and the antibiotic action.

We systematically identified interactions between translation inhibitors and bottlenecks by their deviation from independence. In general, antibiotic action can be alleviated or aggravated by a given bottleneck, *i.e.*, the bacteria can be less or more sensitive to the antibiotic due to the bottleneck, respectively. We quantified the magnitude of these effects by bottleneck dependency (*BD*) scores (Methods) and collected them into a single bottleneck dependency vector per antibiotic. The components of this vector describe the interaction between that antibiotic and all six translation bottlenecks. Bottleneck dependency vectors were diverse (Fig. 3D), indicating that bottlenecks at different stages of the translation cycle differentially affect antibiotic efficacy. These results are consistent with the hypothesis that the high diversity of drug interactions between translation inhibitors (Fig. 1D) originates in the diversity of translation steps targeted by the drugs (Fig. 1A).

The bottleneck dependency vector of a given antibiotic provides a quantitative, functional summary of its interaction with the translation cycle. In this sense, it is a characteristic “fingerprint” of the antibiotic. Clustering of antibiotics based on these bottleneck dependency vectors (Methods) robustly grouped together antibiotics with similar mode of action (CRY and FUS, LCY and CHL in Fig. 3E, respectively). Further, drug interactions between antibiotics from the same cluster were strictly additive (Figs. 1D and 3E). These results show that interactions of antibiotics with translation bottlenecks have considerable explanatory power for drug mode of action and indicate that antibiotics acting as substitutes for one another can be identified based on these interactions.

To challenge the predictive power of translation bottlenecks, we tested whether the mode of action of a partially characterized antibiotic can be inferred from its bottleneck dependency vector. We focused on lamotrigine (LAM), an anticonvulsant drug which was recently identified to inhibit maturation and in turn reduce the number of translating ribosomes, potentially by interfering with initiation factor 2 (IF2, encoded by *infB*) [Stokes *et al.*, 2014]. The bottleneck dependency vector of LAM was most similar to that of KSG (Fig. 3D,E). As for LAM, a reduction of translating ribosomes is a signature of the initiation inhibitor KSG [Kaberdina *et al.*, 2009]. Hence, this observation further corroborates that the similar bottleneck dependency vectors for translation inhibitors indicate similar mode of action.

We further tested how an antibiotic with a mode of action unrelated to translation interacts with translation bottlenecks. If drug interactions are primarily determined by their mode of action [Yeh *et al.*, 2006; Brochado *et al.*, 2018], antibiotics interfering with processes unrelated to translation should be affected similarly by all different translation bottlenecks as the net effects of translation bottlenecks are indistinguishable – all lead to cessation of protein synthesis. To test this idea, we chose the antibiotic trimethoprim (TMP), which inhibits folate synthesis by binding to dihydrofolate reductase and is not known to directly perturb translation [Walsh, 2003]. Its bottleneck dependency vector indicates that all bottlenecks alleviated TMP’s action to various degrees (Fig. 3D) – a characteristic that is incompatible with any of the clusters of translation inhibitors (Fig. 3E). Furthermore, TMP is known to primarily interact antagonistically or suppressively with translation inhibitors [Bollenbach *et al.*, 2009; Yeh *et al.*, 2006]. These results support the idea that the effects of specific translation bottlenecks are diverse for antibiotics targeting translation, but not for antibiotics with modes of action unrelated to translation.

Streptomycin stands out among translation inhibitors, as its action is aggravated by all translation bottlenecks (Fig. 3D). This might be a consequence of additional unspecific modes of action. We corroborated this by measuring the bottleneck dependency vector of a prodrug nitrofurantoin (NIT). Nitrofurantoin has complicated effects on the bacterial cell, including the formation of non-native disulfide bonds in protein structures [Bandow *et al.*, 2003], DNA damage, and oxidative stress [Mitosch *et al.*, 2017]. A similar bottleneck dependency between STR and NIT likely reflects that, beyond inhibiting translation, STR has strong secondary effects: it causes protein mistranslation, changes in membrane potential, and membrane permeabilization [Davis, 1987]. Some of these processes, in particular the production of dysfunctional proteins, overlap with those of NIT [Bandow *et al.*, 2003], offering an explanation for the observed similarity of these seemingly unrelated drugs.

### 2.4 Drug interactions can be predicted from antibiotic responses to translation bottlenecks

We reasoned that the effects of translation bottlenecks on antibiotic action should also have predictive power for drug interactions involving translation inhibitors. We thus sought for a quantitative way of probing the contribution of translation bottlenecks to drug interactions between translation inhibitors. Translation can be seen as a sequence of steps in which ribosomes progress through the protein production cycle. Antibiotics and genetic translation bottlenecks hinder this progression similarly by reducing the transition rates between such steps (Fig. 4A). In cases where an antibiotic specifically targets a single translation step and reduces the same transition rate as a genetic translation bottleneck, the antibiotic effect and the genetic translation bottleneck should be equivalent perturbations, *i.e.*, the consequences of any perturbation elsewhere in the translation cycle should be independent of the exact means by which such a reduction has been effected (Fig. 4B).

**Figure 4:**
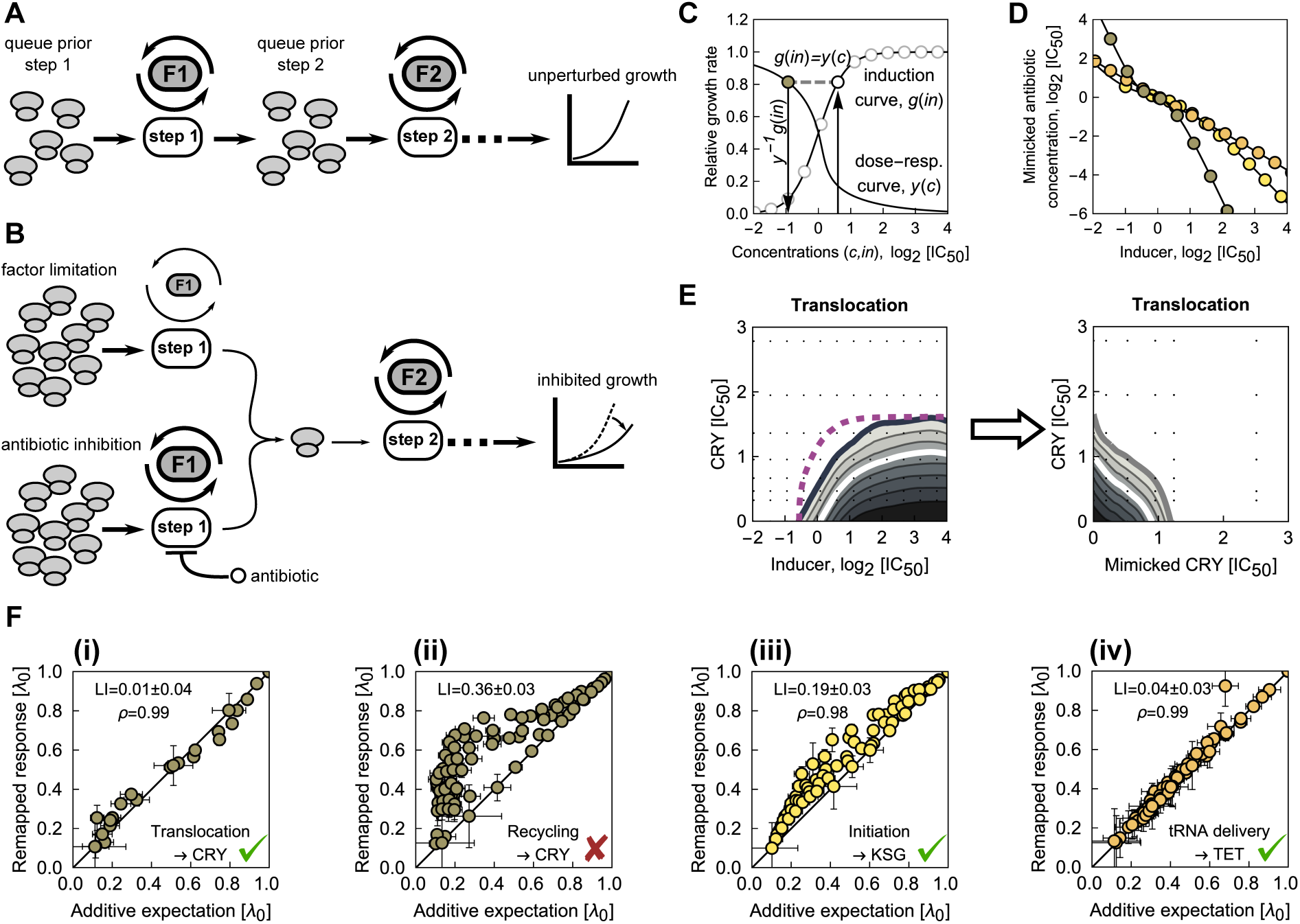
Translation factor deprivation mimics the action of equivalent antibiotics. **(A)** Schematic of translation as a sequence of steps (white), catalyzed by translation factors (gray). In the absence of perturbations, ribosomes progress through the steps unimpeded, resulting in unperturbed growth. **(B)** Schematic of perturbed translation. Top: as the abundance of factor F1 is lowered (smaller factor symbol), the rate of step 1 decreases (thinner arrows) and ribosomes queue in front of the bottleneck. Bottom: the same rate is reduced by an antibiotic. The effects of factor deprivation and antibiotic action on growth are equivalent. **(C)** Schematic of conversion of inducer concentration *in* (here for the translocation factor) into the mimicked antibiotic concentration *c* (here: CRY). For each inducer concentration *in*, the growth rate from the induction curve *g*(*in*) is determined and the same growth rate on the antibiotic dose-response curve *y* (*c*) is identified (gray dashed line); the inverse function of the dose-response curve yields the equivalent antibiotic concentration as *c* = *y* ^−1^ (*g*(*in*)). **(D)** Resulting conversion of inducer concentration *c*_*i*_ into antibiotic concentration *c* for three different pairs of equivalent perturbations: CRY-translocation (gray), KSG-initiation (yellow) and TET-tRNA delivery (orange). **(E)** Inducer-antibiotic response surface (left) and mimicked antibiotic-antibiotic response surface (right) upon conversion of inducer concentration as in (C) and (D). Purple dashed line shows isobole for multiplicative responses at relative growth rate 0.2. The remapped response surface is additive, corroborating the equivalence of CRY and translocation factor deprivation. **(F)** Comparison of response surfaces remapped as in E to the additive expectation. The bottlenecks and antibiotics are shown on the bottom right, respectively. Errors in *LI* and in expected and remapped responses were evaluated by bootstrapping (Methods). **(i)** Example from (E): additive expectation and remapped response surface agree (*ρ* = 0.99). **(ii)** As (i), but for a recycling bottleneck. The large and statistically significant discrepancy in *LI* from 0 indicates that CRY and recycling bottleneck are not equivalent (Methods, Fig. S4). **(iii)** As (i), but for KSG and initiation bottleneck (*ρ* = 0.98). **(iv)** As (i), but for TET and tRNA delivery bottleneck (*ρ* = 0.99).

To establish the equivalence between translation bottlenecks and antibiotic action, we first transformed the measurements of growth rate as a function of translation factor induction into dose-response curves of a corresponding idealized antibiotic that targets a single translation step with perfect specificity. In essence, this procedure converts inducer concentrations into equivalent antibiotic concentrations: the two concentrations are identified as equivalent if they lead to the same relative growth rate (Fig. 4C,D; Methods). If the perturbations of factor and antibiotic are equivalent, then the true and idealized antibiotic should act as substitutes for each other, and thus exhibit an additive drug interaction. Consequently, we can use this comparison (Figs. 4C and S4) to test systematically if the action of antibiotics is equivalent to specific translation bottlenecks.

We found that the effect of certain translation inhibitors can be almost perfectly mimicked by translation bottlenecks. Within our selection of antibiotics, several strong candidates for equivalent perturbations exist (Fig. 1A): CRY, FUS and STR with EF-G (translocation); KSG with IF2 (initiation); and TET with EF-Tu (tRNA-delivery). For example, remapping the response surface of CRY and EF-G yields an additive surface (Fig. 4E), corroborating that CRY and the EF-G translocation bottleneck are equivalent perturbations. In contrast, if the bottleneck is not equivalent to the drug, remapping does not yield an additive response surface; an example is CRY and the recycling bottleneck (Fig. 4F,ii). In general, demonstrating that an antibiotic acts as an equivalent perturbation to a specific translation factor provides strong evidence for its primary mode of action, since translation factors are thought to control individual steps with high specificity.

For antibiotics that are equivalent to specific translation factors (Fig. 4F), drug interactions with other antibiotics can be directly explained and predicted. In practice, this is done by remapping the antibiotic-translation factor response surfaces as described above (Fig. 5A,B). The resulting prediction will be faithful if the drug interaction originates exclusively from the combination of two bottlenecks in the translation cycle. Drug interactions predicted using this procedure were often highly accurate (Fig. 5C). In particular, some of the most striking cases of antagonistic and suppressive interactions were correctly predicted. For example, the suppressive interaction of CHL with FUS was correctly predicted, including its direction: FUS loses potency when exposed to CHL (Fig. 5C-i). Further, the prediction of antagonism between CHL and STR was qualitatively correct (Fig. 5C-ii). Similarly, prediction of these interactions with FUS and STR were also correct for LCY (Fig. S5) which is similar to CHL (Fig. 3E). The remapping approach further correctly predicted the prevalent antagonism and suppression of the initiation inhibitor KSG with other translation inhibitors (Fig. 1D). Remapping qualitatively accounted for all observed interactions of KSG with quantitative agreement in several cases, including KSG-CHL (Fig. 5A-ii) and KSG-STR (Fig. 5C-iv and SI). Thus, several drug interactions with previously elusive mechanisms are explained by the interplay of the specific steps in the translation cycle that are targeted by the constituent antibiotics.

**Figure 5:**
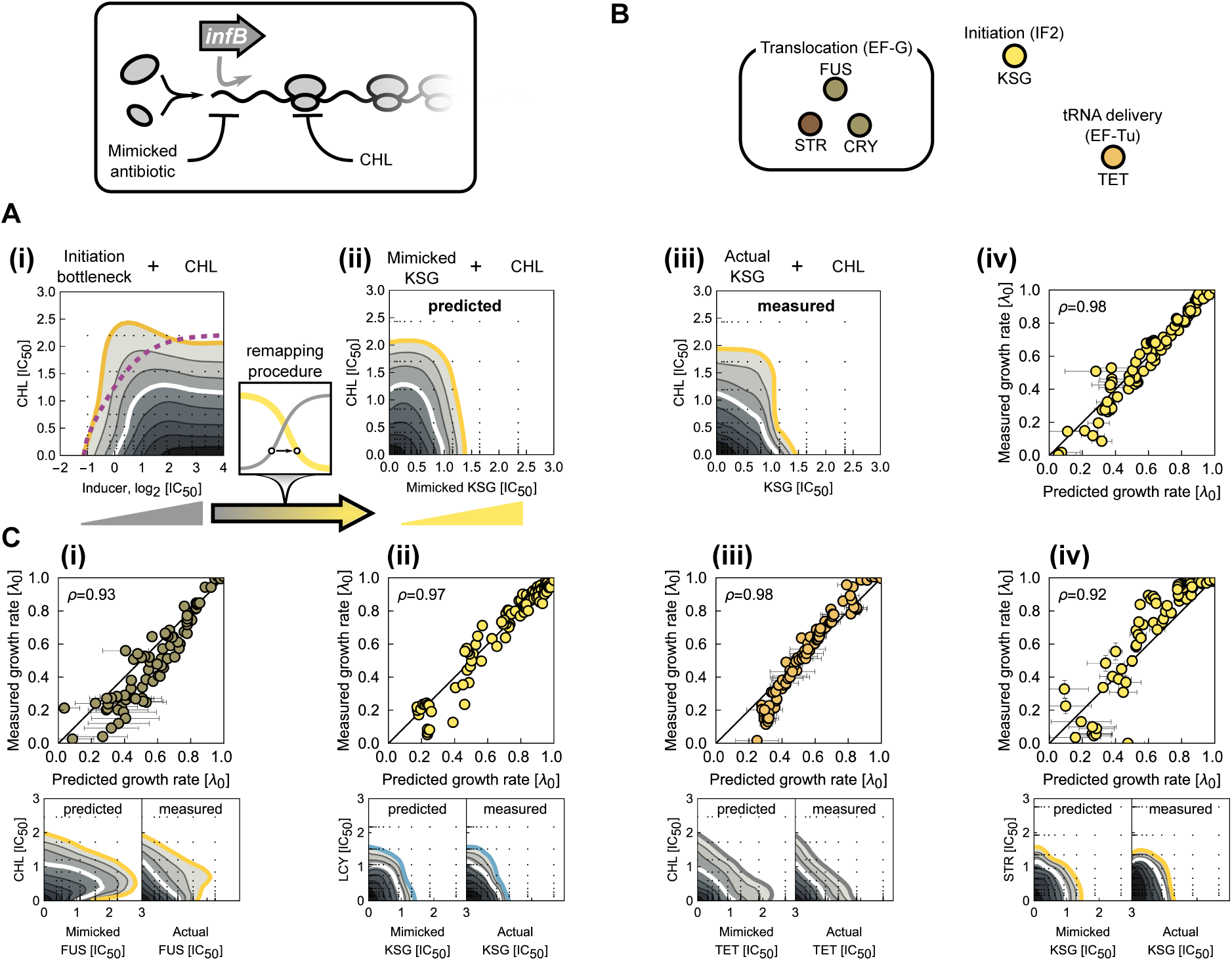
Equivalent translation bottlenecks can predict antibiotic interactions. **(A)** Example of drug interaction prediction based on equivalent translation bottlenecks. **(i)** Response surface of CHL combined with the inducer for the initiation (*infB*) bottleneck shows mild alleviation. This response surface contains information about the interaction between CHL and any antibiotic that interferes with initiation. The inducer axis is remapped into mimicked antibiotic concentration (lower box; Fig. 4C-E). **(ii)** Resultant prediction of response surface for the initiation-inhibiting antibiotic KSG and CHL. **(iii)** Measured KSG-CHL response surface for direct comparison; strong antagonism is observed as predicted. **(iv)** Point-by-point comparison of predicted and measured response surfaces (Pearson’s *ρ* = 0.98). **(B)** Schematic showing antibiotics and their equivalent translation factor bottlenecks. Drug interactions with these antibiotics can be predicted for any antibiotic with known response to the equivalent bottleneck. Color-code shows cluster identity from Fig. 3E. **(C)** Comparison of predicted and measured response surfaces for different antibiotics in combination with antibiotics that have a factor analog. Top row: scatter plots as in A-iv; bottom row: predicted and measured response surfaces, respectively. **(i)** Suppression of FUS by CHL at high inhibition is correctly predicted. **(ii)** Antagonistic interaction between KSG-LCY (KSG is mimicked by initiation bottleneck) is correctly predicted. **(iii)** Additivity between CHL-TET based on mimicking TET by a tRNA delivery bottleneck is correctly predicted. **(iv)** Strong antagonism between KSG and STR based on mimicking KSG by an initiation bottleneck is correctly predicted.

Our approach further explained nontrivial additive interactions. In particular, the additive interaction between CHL and TET is hard to rationalize: these antibiotics have completely different binding sites on the ribosome. However, CHL and TET interacted similarly with translation bottlenecks (Fig. 3E) and their interaction was faithfully captured by the remapping approach (Fig. 5C-iii). This observation suggests that the action of CHL is largely equivalent to inhibiting tRNA delivery. As CHL binding interferes with a distal end of tRNA on the A-site [Wilson, 2014], this suggests that perturbation of tRNA dynamics is at the heart of the drug interaction between TET and CHL. KSG and ERM constitute another antibiotic pair that interacted additively and was clustered together. Remapping correctly predicted additivity between KSG-ERM (SI); however, ERM does not directly inhibit initiation as does KSG (Table 1). Yet, it is likely that the inability of ERM to inhibit translation when the nascent peptide chain is extended beyond a certain length effectively leads to a functional equivalence, which results in additivity and co-clustering of ERM and KSG.

For certain antibiotic pairs, the predictions based on equivalent translation bottlenecks failed to explain the observed drug interactions (*e.g.*, for LCY-CRY and CHL-CRY; SI), indicating that these interactions have origins outside of the translation cycle. We expect that these cases are often due to idiosyncrasies of the drugs, which will require separate in depth characterization in each case. In contrast, our results show that various non-trivial drug interactions between antibiotics are systematically explained by the interplay of specific bottlenecks in the translation cycle that are caused by the antibiotics. While the growth-law based biophysical model already explained ≈57% (16 of 28) of the observed interactions (Fig. S2), suppressive interactions were only captured after taking into account the multi-step nature of translation (Fig. S5), thus increasing the explained fraction to ≈71% (20 of 28). If suppressive drug interactions are caused by the interplay of different translation bottlenecks alone, it should be possible to recapitulate these interactions in a purely genetic way. We thus expanded our approach of using genetic translation bottlenecks as proxies for antibiotics by introducing multiple genetic bottlenecks simultaneously in the same cell.

### 2.5 Simultaneous titration of translation factors reveals robust suppression between translocation and initiation inhibition

We focused on the interactions between initiation inhibitors (such as KSG) and translocation inhibitors (such as CRY, STR, FUS) as they were exclusively antagonistic or suppressive (Fig. 1D). Moreover, the initiation inhibitor KSG alleviated a genetic translocation bottleneck and an initiation bottleneck in turn suppressed the effect of the translocation inhibitor FUS (Fig. 3C). These observations suggest that a universal mechanism underlies the suppression between initiation and translocation inhibitors.

Thus, we constructed a synthetic strain that enables simultaneous independent control of initiation and translocation factor levels. We integrated the initiation and translocation factors outside their native loci under tight control of promoters inducible by IPTG and anhydrotetracycline (aTc), respectively, in a strain in which their endogenous copies were deleted (Figs. 6A and S6; Methods). To maximize the precision of induction that is achievable with different inducer concentrations, we put both factors under negative autoregulatory control by chromosomally integrated repressors [Klumpp *et al.*, 2009; Scott *et al.*, 2010]. The resulting strain showed no growth when at least one of the inducers was absent but wild type growth was fully rescued in the presence of both inducers (Fig. 6B). These observations confirm that both translation factors are essential and show that their expression can be varied over the entire physiologically relevant dynamic range, thus enabling quantitative genetic control of two key translation processes.

**Figure 6:**
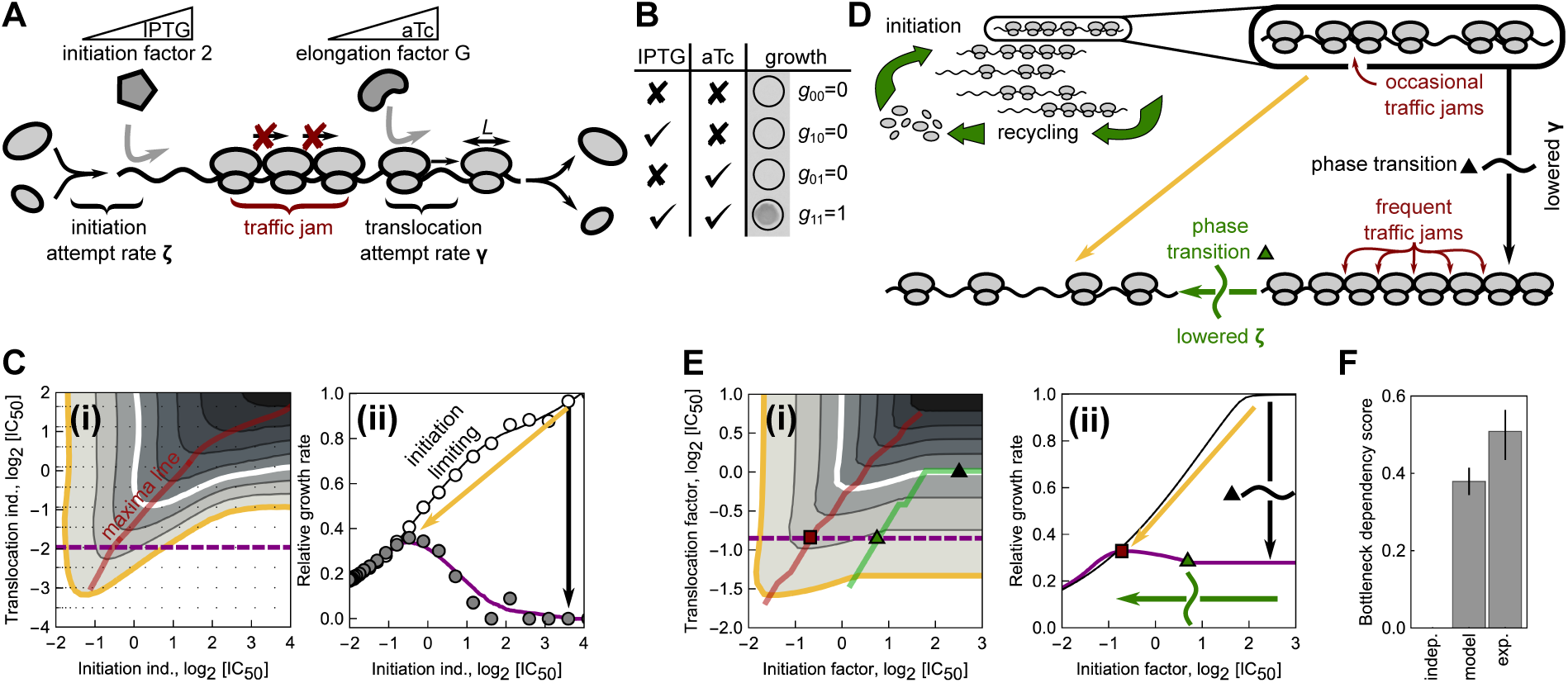
Suppression between inhibition of translocation and initiation is explained by dissolution of ribosome traffic jams in a phase transition. **(A)** Schematic of ribosomes progressing along a transcript – a stuck ribosome can cause a traffic jam. Ribosomes undergo factor-mediated initiation events with attempt rate *ζ* and translocation with attempt rate *γ*. Expression of initiation and elongation factor G (translocation) are controlled by level of inducer (IPTG and aTc, respectively). **(B)** Results of all-or-nothing growth assay: bacteria grow only when both essential factors are induced. **(C;i)** Measured growth rate response surface for the dual inducible promoter strain from (A) as a function of both inducer concentrations; red line shows ridge of maximum growth. **(C;ii)** Cross-section of the response surface along dashed purple line in (i) (gray circles) and at maximal aTc induction (white circles); solid lines are smoothed profiles. Black arrow denotes decrease in translocation; if initiation is lowered simultaneously with translocation (orange arrow), growth reduction is smaller. **(D)** Schematic of theoretical model: translation is described as an ensemble of transcripts competing for the limited and growth-rate-dependent pool of ribosomes. Ribosomes advance on transcripts as described by a generalized totally asymmetric simple exclusion process (TASEP) for particles of size *L*, see (A) and text. When *γ* < *ζ*(1 + *L*^1/2^), ribosomes saturate and traffic jams develop, resulting in a drop in elongation and growth (black arrow, transition happens at black triangle) (Methods, [Klumpp and Hwa, 2008; Lakatos and Chou, 2003]). When *ζ* < *γ*/(1 + *L*^1/2^), a phase transition occurs (green triangle): traffic jams dissolve; elongation and growth increase (along the green arrow). **(E;i)** Growth rate predicted by the generalized TASEP model recapitulates suppression of translocation inhibition by lowered initiation; note that, unlike in (C), axes show the concentrations of translation factors. States below and to the right of the green line are in the translocation limiting regime. **(E;ii)** Cross-sections of the response surface: solid purple line corresponds to dashed purple line in (i). As the initiation factor level is decreased, the critical point of the phase transition (green triangle) is reached; growth starts increasing after passing the critical point, and decreases again after passing the maximum (red square) as the number of translating ribosomes becomes limiting. **(F)** Bottleneck dependency (*BD*) score quantifies the deviation from independent expectation (*BD* = 0) for the response surfaces in (C;i) and (E;i); error bars are 5%-95% bootstrap intervals.

Curtailing translation initiation suppresses the effect of a genetic translocation bottleneck. We determined the bacterial response to varying translocation and initiation factor levels by measuring growth rates over finely resolved two-dimensional concentration gradients of both inducers. The resulting response surface clearly showed that inhibition of initiation alleviates the effect of translocation inhibition (Figs. 6C and S6). This phenomenon exactly mirrors the antibiotic-antibiotic (KSG-FUS, Fig. 1D) and bottleneck-antibiotic interactions (initiation-FUS, Fig. 3C). Note that an all-or-nothing approach (Fig. 6B), which is analogous to common genetic epistasis measurements [Constanzo *et al.*, 2010], would miss this suppressive effect, highlighting the importance of the quantitatively controlled perturbations we used. Taken together, these data show that the interplay of translation initiation and translocation alone is sufficient to produce strong suppression: dialing down initiation cranks up growth stalled by translocation bottlenecks. The widespread suppression between antibiotics targeting initiation and translocation is thus explained as a general consequence of the combined inhibition of specific translation steps alone.

What is the underlying mechanism of the suppressive interaction between initiation and translocation inhibitors? We hypothesized that this suppression results from alleviating ribosome “traffic jams” that occur during translation of transcripts when the translocation rate is low (Fig. 6D). The traffic of translating ribosomes that move along mRNAs can be dense [Mitarai *et al.*, 2008] and when a ribosome gets stuck (*e.g.*, due to a low translocation rate), it blocks the translocation of subsequent ribosomes. The resulting situation is similar to a traffic jam of cars on a road. Traffic jams form due to asynchronous movement and stochastic progression of particles in discrete jumps, which is a good approximation for the molecular dynamics of a translating ribosome. If particle progression were deterministic and synchronous, no traffic jams would form. A classic model of queued traffic progression, which can be applied to protein translation [MacDonald *et al.*, 1968; MacDonald and Gibbs, 1969], is the Totally Asymmetric Simple Exclusion Process (TASEP) [Shaw *et al.*, 2003; Zia *et al.*, 2011].

We developed a variant of the TASEP that describes the traffic of translating ribosomes on mRNAs and takes into account the laws of bacterial cell physiology. There are several differences between the classic TASEP and translating ribosomes moving along a transcript. First, a ribosome does not merely occupy a single site (codon), but rather extends over 16 codons [Kang and Cantor, 1985]. Second, the total number of ribosomes in the cell is finite and varies as dictated by bacterial growth laws [Scott *et al.*, 2010; Scott *et al.*, 2014]. Third, translation steps are mediated by translation factors that bind to the ribosome in a specific state and push the ribosome into another state [Rodnina, 2018]. These transitions are stochastic with rates that depend on the abundance of ribosomes in a specific state and on the abundance of translation factors available to catalyze the step. Thus, the initiation and translocation-attempt rates, which are constants in the classic TASEP, depend on the state of the system. We formulated a generalized TASEP that captures these extensions, estimated all of its parameters based on literature, and derived the model equations analytically (Methods); the resulting growth rate was calculated numerically. In brief, our generalized TASEP model provides a physiologically-realistic description of the factor-mediated traffic of ribosomes on multiple transcripts.

Without any free parameters, the generalized TASEP qualitatively reproduced the suppressive effect of lowering the initiation rate under a translocation bottleneck (Fig. 6E). This suppression results from a phase transition between the translocation- and the initiation-limited regime (Methods). In the translocation-limited regime (black arrow in Fig. 6E-ii), ribosome traffic is dense and cannot be further increased by boosting the initiation attempt rate. Upon decreasing the initiation attempt rate *α* (green arrow in Fig. 6E-ii), a phase transition to the initiation-limited regime occurs. Beyond the critical point of this phase transition (green triangle in Fig. 6E-ii), the elongation velocity, and with it the growth rate, begins to increase with decreasing initiation attempt rate. Thus, ultimately, a non-equilibrium phase transition in which ribosome traffic jams dissolve underlies the suppressive effect. To compare measured and predicted surfaces that have different axes, we calculated their deviation from independence as for the bottleneck dependency score (Figs. 3D and S3). By this measure, the model faithfully captured the clear deviation from the multiplicative expectation (Fig. 6F); the agreement with the experimental data is surprisingly good, especially since the model results are parameter-free and not a fit to the experimental data.

Taken together, these results show that suppressive drug interactions between translation inhibitors are caused by the interplay of two different translation bottlenecks. Close agreement of the experiments with a plausible theoretical model of ribosome traffic, which captures physiological feedback mediated by growth laws, strongly suggests that suppression is caused by ribosome traffic jams. Such traffic jams result from imbalances between translation initiation and translocation; they dissolve in a phase transition that occurs when one of these processes is slowed, leading to an overall acceleration of translation and growth. Thus, a non-equilibrium phase transition in ribosome traffic is at the heart of suppressive drug interactions between antibiotics targeting translation initiation and translocation.

## 3 Discussion

We established a framework that combines mathematical modeling, high-throughput growth rate measurements, and genetic perturbations to elucidate the underlying mechanisms of drug interactions between antibiotics inhibiting translation. Kinetics of antibiotic-target binding and transport together with the “growth laws”, *i.e.*, bacterial response to translation inhibition (Fig. 2), form a biophysically-realistic baseline model for predicting antibiotic interactions from properties of individual antibiotics alone. This model explained some interactions, but not all, failing specifically on suppressive interactions. Predictions improved by taking into account the step-wise progression of ribosomes through the translation cycle (Fig. 4, 5). This was achieved by mimicking antibiotic perturbations of this progression genetically, which directly identified the contribution of antibiotic-imposed translation bottlenecks to the observed drug interactions. Finally, to explain the origin of suppressive interactions unaccounted for by the biophysical model, we modeled the traffic of translating ribosomes explicitly. Our results show that translocation inhibition can cause ribosomal traffic jams, which dissolve in a non-equilibrium phase transition when initiation is inhibited simultaneously with translocation, thereby restoring growth (Fig. 6). This phase transition explains the suppressive drug interactions between antibiotics targeting initiation and translocation.

Taken together, our framework mechanistically explains twenty out of twenty-eight observed drug interactions (Fig. 1, S2, S5), as judged by highly stringent quantitative and statistical criteria (Methods). Here, even the cases rejected as quantitatively different are insightful. For example, the remapping-based prediction of CHL-FUS interaction (Fig. 5C-ii) over-estimates the suppression and is rejected on quantitative basis; yet remapping correctly suggests the occurrence of suppression and its direction. Qualitative observations like these deepen our understanding of drug interactions as they highlight potential mechanisms of drug interaction, on top of which additional mechanisms are acting. While we focused on translation inhibitors, key elements of our framework can be generalized to drugs with other modes of action. Specifically, when considering a drug that targets a specific process mediated by an essential enzyme, our method of equating the deprivation of the enzyme with the action of an antibiotic is readily applicable.

Mimicking the effects of two drugs with controllable genetic perturbations generalizes the concept of genetic epistasis to continuous perturbations. Epistasis studies compare the effects of double gene knockouts to those of single knockouts and identify epistatic interactions – an approach that can reveal functional modules in the cell [Segre *et al.*, 2005; Tong *et al.*, 2004; Constanzo *et al.*, 2010]. Our results show that continuous genetic perturbations provide essential additional information on genetic interactions (Fig. 6). Firstly, the direction of epistatic interactions cannot be extracted from measurements of single and double mutants. Secondly, the quantitative information obtained from such “continuous epistasis” measurements provides more stringent constraints for mathematical models of biological systems. In particular, continuous epistasis data can be powerful for the development of whole-cell models that describe the interplay of different functional modules in the cell. Thirdly, this approach allows including essential genes in epistatic interaction networks even for haploid organisms, which otherwise requires the use of less well-defined hypomorphs. Thus, continuous epistasis measurements as in Fig. 6C augment all-or-nothing genetic perturbations.

Continuous epistasis measurements further enable a deeper understanding of previously mysterious antibiotic resistance mutations. Specifically, translation bottlenecks that alleviate the effect of an antibiotic expose a latent potential for resistance development. Indeed, mutations leading to effects that are equivalent to factor-imposed bottlenecks occur under antibiotic selection pressure. For example, resistance to ERM in *E. coli* can be conferred by mutations in proteins of the large ribosomal subunit, which hinder its maturation and lower its stability [Zaman *et al.*, 2007]. Consistent with this observation, our results indicate that the action of ERM is alleviated by lowering the stability of the 50S subunit (Fig. 3D). Furthermore, mutations in recycling factor were selected in *Pseudomonas aeruginosa* evolved for resistance to the TET derivative tigecycline [Sanz-Garcia *et al.*, 2018]. The observed alleviation of TET action by a recycling bottleneck (Fig. 3D) offers a mechanistic explanation for the beneficial effects of these mutations. Mutations in other genes predicted based on the effect of translation bottlenecks may be difficult to observe, especially in clinical isolates, due to the associated fitness cost and selection pressure for reverting the mutations in the absence of the antibiotic. Beyond mutations conferring resistance to individual drugs, consistent or conflicting dependencies of different antibiotics on translation bottlenecks may further indicate the potential for evolving cross-resistance and collateral sensitivity, respectively [Baym *et al.*, 2016].

In conclusion, we presented a systematic approach for discovering the mechanistic origins of drug interactions between antibiotics targeting translation. As the translation machinery is highly conserved, the interaction mechanisms for drugs targeting specific steps of translation we uncovered may generalize to diverse other organisms. Our approach of mimicking drug effects with continuous genetic perturbations is general and can be extended to antibiotics with other primary targets, other types of drugs, and other organisms. Our quantitative analysis relies on the established correlation between ribosome content and growth rate in varying nutrient environments [Scott *et al.*, 2010]. This highlights the importance of elucidating such growth laws in other organisms for gaining a deeper understanding of combined drug action. In the long run, extending our combined experimental-theoretical approach to other types of drugs and other biological systems will enhance our understanding of drug modes of action and interaction mechanisms and provide deeper insights into cell physiology.

## 4 Acknowledgments

We thank M. Hennessey-Wesen, I. Tomanek, K. Jain, A. Staron, K. Tomasek, M. Scott, and Z. Gitai for reading the manuscript and constructive comments. K.B. is indebted to C. Guet for additional guidance and generous support which rendered this work possible. K.B. thanks all members of Guet group for many helpful discussions and sharing of resources. K.B. additionally acknowledges the tremendous support from A. Angermayr and K. Mitosch with experimental work. We further thank E. Brown for helpful comments regarding lamotrigine. This work was supported in part by Austrian Science Fund (FWF) standalone grants P 27201-B22 (to T.B.) and P 28844 (to G.T.), HFSP program Grant RGP0042/2013 (to T.B.), and German Research Foundation (DFG) Collaborative Research Centre (SFB) 1310 (to T.B.).

## 5 Author contributions

Conceptualization, methodology, formal analysis, investigation, writing – original draft, writing – review & editing: K.B., G.T., and T.B. Funding acquisition, resources, and supervision: G.T., and T.B.

## 6 Declaration of Interests

The authors declare no competing interests.

## 7 Material and Methods

### Bacterial strains

*Escherichia coli* K-12 MG1655 strain was used as wild-type (WT) strain. When necessary, selection on kanamycin was performed at 25 *µ*g/mL (for post-recombineering selection, see below) or at 50 *µ*g/mL (for P1 transduction and plasmid selection). A concentration of 100 *µ*g/mL was used for ampicillin (pCP20, resistance cassette resolution) and spectinomycin (pSIM19, recombineering).

To measure the bioluminescence time traces, pCS-*λ* encoding the bacterial *luxCDABE* operon driven by the constitutive *λ*-P_R_ promoter was transformed into the strains of interest [Kishony and Leibler, 2003]. Selection for the luminescence plasmid was used during the preparation of glycerol stocks (kanamycin 50*µ*g/mL) but was omitted during the measurements to avoid unknown chemical interactions between used antibiotics. The plasmid was stably maintained as we observed no significant fitness deficit due to pCS-*λ* and no apparent spontaneous loss of the plasmid.

The translation factor titration platform was established in a strain HG105 (MG1655 *ΔlacIZYA*) [Garcia *et al.*, 2011]. Briefly, endogenous genes encoding for translation factors were first sub-cloned into pKD13 vector under the control of P_LlacO–1_ promoter with FRT-flanked kanamycin resistance cassette (kan^R^) and *rrnB* terminator *TrrnB* upstream and downstream of the gene, respectively [Datsenko and Wanner, 2000; Klumpp *et al.*, 2009; Scott *et al.*, 2010; Lutz and Bujard, 1997]. The tandem of kan^R^ and a gene with all regulatory elements was integrated into the chromosome (*galK* locus) using *λ*-red recombineering (plasmid pSIM19 [Datta *et al.*, 2006]). The kanamycin resistance cassettes here and in the following steps were resolved using yeast FLP resolvase expressed from pCP20 [Cherepanov and Wackernagel, 1995]. Loss of the resistance cassette and curing of the pCP20 plasmid were checked by streaking on selection agar plates with antibiotics and by junction PCR (for resolution). Following the resolution of kan^R^, the endogenous factor was inactivated by in-frame deletion: kan^R^ was integrated into the gene locus and then resolved, which left a 34 aa-residue peptide [Datsenko and Wanner, 2000]. We were unable to introduce kan^R^ directly into the strain with P_LlacO–1_ driven *frr*; therefore, we first performed the deletion in an auxiliary strain MG1655 *Δfrr*::kan^R^ bearing the ASKA plasmid with *frr* [Kitagawa *et al.*, 2005] [JW0167(-GFP)], which complemented the chromosomal deletion when IPTG was added. Deletion was possible in the auxiliary strain. We then moved the deletion by generalized P1 transduction [Lennox, 1955]. For *tufAB*, we P1-transduced the deletions (*ΔtufA*::kan^R^ and *ΔtufB*::kan^R^) sequentially from the respective gene deletion strains from the KEIO collection [Baba *et al.*, 2006]. All other deletions were performed directly in the strains of interest using *λ*-red recombineering using pKD13 as a template for the cassette amplification [Datsenko and Wanner, 2000]. In the last step, *lacI* driven by the P_LlacO–1_ promoter (yielding a growth-rate independent negative autoregulation [Klumpp *et al.*, 2009; Scott *et al.*, 2010]) together with the FRT-flanked kan^R^ was integrated into *intS* locus and the resistance cassette was resolved. The allele *ΔintS*::kan^R^-P_LlacO–1_*-lacI-TrrnB* was moved into the strains by generalized P1 transduction. All chromosomal modifications were validated by PCR. The factor titration platform and the repressor operon were Sanger-sequenced at the integration junctions using PCR primers or a primer binding into the kan^R^ promoter region (which is upstream of the P_LlacO–1_ promoter prior the resolution). The final genotype for the strains bearing the factor titration platforms is HG105 *ΔgalK*::frt-P_LlacO–1_-*x Δx*::frt *ΔintS*::frt-P_LlacO–1_-*lacI*, where *x* denotes the chosen factor. These strains contained no plasmids and no antibiotic resistance cassettes but had a single copy of a translation factor under inducible control.

To generate the strain with independently regulated initiation and translocation factors, we started with a strain carrying a single *infB* copy driven by P_LlacO–1_. Then, the negatively autoregulated *tetR* repressor was integrated into the chromosome, followed by FLP resolvase-mediated resolution of the selection marker. This enabled the integration of P_LtetO–1_-driven *fusA* into the *intS* locus; resolution was followed by the disruption of the endogenous copy of *fusA*. Furthermore, we introduced a negatively auto-regulated *lacI* into the *xylB* locus. This yielded a marker-less strain with the two essential genes *infB* and *fusA* under inducible, negatively autoregulated and independent control. The final genotype is: HG105 *ΔgalK*::frt-P_LlacO–1_-*infB ΔinfB*::frt *ΔycaCD*::frt-P_LtetO–1_-*tetR ΔintS*::frt-P_LtetO–1_-*fusA ΔfusA*::frt *ΔxylB*::frt-P_LlacO–1_-*lacI*. Oligonucleotide sequences, targeted template, restrictions sites (when used) and brief description of use are in Supplementary Table S2. All DNA modifying enzymes and Q5 polymerase used in PCR were from New England Biolabs.

### Growth rate assay and two-dimensional concentration matrices

Rich lysogeny broth (LB) medium, which at 37°C supports a growth rate of 2.0 ± 0.1 h^−1^, was used. LB medium was prepared from Sigma Aldrich LB broth powder (L3022), pH-adjusted by adding NaOH or HCl to 7.0 and autoclaved. Antibiotic stock solutions were prepared from powder stocks (for catalog numbers, see Table S1), dissolved either in ethanol (CHL, ERM and TET), DMSO (LAM and TMP) or water (KAN, CRY, LCY, KSG, FUS and STR), 0.22 *µ*m filter sterilized and kept at −20°C in the dark until used. Antibiotics were purchased from Sigma Aldrich or AvaChem. A previously established growth-rate assay based on photon counting was used to precisely quantify the absolute growth rates over the course of 5-9 generations [Kishony and Leibler, 2003]. Cultures were grown in 150 *µ*L of media in white 96-well microtiter plates (Nunc 236105), which were tightly sealed by transparent adhesive foils (Perkin-Elmer 6050185 TopSeal-A PLUS) to prevent contamination and evaporation. We prepared glycerol stocks of WT and factor-titration platform strains from saturated overnight cultures. We inoculated the cultures with ∼ 10^2^ cells per well (1:10^6^ dilution) from either thawed glycerol stocks (for the drug interaction network) or from liquid cultures in which we first incubated the bacteria containing the factor titration-platform for 1 h in the absence of IPTG (inoculated by 1:2000 dilution of the glycerol stock) to partially dilute out the remaining factor molecules before additional 1:1000 dilution into measurement plates. Between 10-20 plates were cycled through a plate reader using a stacking system (Tecan M1000). We built a custom incubator box around the stacker towers to facilitate ventilation and fix the temperature to 37°C. This incubator was designed and troubleshot by BK and Andreas Angermayr (IST Austria and University of Cologne) and built by IST Miba Machine Shop. Each plate was read every 20-40 min and was shaken (orbital 10 s, 582 rpm) immediately before reading (settle time10 ms, integration time 1 s). Plates were manually pipetted and concentration gradients of antibiotics and IPTG were prepared by serial dilution (0.70-fold). Growth rates were determined as a best-fit slope of a linear function fitted to the log-transformed photon counts per second. The detailed fitting procedure and examples of growth curves are shown in Fig. S1. The experimental and analysis procedure led to reproducible measurements of growth rates between days (Fig. S1, *ρ* ≈ 0.86). Two-dimensional gradients were usually set up in a 12×16 matrix (across two 96-well plates). For the double factor titration experiment the inducer gradients were set up across 6 plates to form a 24×24 grid.

### Normalization of dose-response surfaces

All growth rates were normalized with respect to the average growth rate in drug-free medium [for factor-titration strains at highest inducer concentration (5 mM)]. Small differences between individual dose-response curves were inevitable due to challenges at preparing identical concentrations gradients on different days. To correct for such day-to-day variability, we rescaled the concentration units to the IC_50_ for each drug. The IC_50_ was obtained from fitting the Hill function *y* (*x*) = 1/ [1 + (*x* /IC_50_)^*n*^] to the individual dose-response curves. The dose-response curve of each drug was measured seven times and averaged. The IC_50_ and corresponding errors reported in Table 1 are extracted from such average dose-response curves. Induction curves were normalized slightly differently, using a shifted and increasing Hill function in the form *g*(*in*) = [(*in* + *in*_0_)/IC_50_]^*n*^ / {1 + [(*in* + *in*_0_)/IC_50_]^*n*^}, where *in*_0_ is a concentration offset. The latter parameter was required as the complete cessation of growth was not achievable in some cases even in the absence of inducer as the promoter P_LlacO–1_ is leaky. Inducer concentrations were thus rescaled via *in* ⟶ (*in* + *in*_0_)/IC_50_.

### Smoothing of dose-response surfaces

To reduce noise when plotting response-surfaces, we smoothed the data using a custom Mathematica script that implements locally weighted regression (LOESS) [Cleveland and Devlin, 1988]. This approach only smoothed the contours and did not alter the character of dose-response surfaces. Smoothing was only used for plotting and not for the analysis in which only linear interpolations between points were used (Mathematica function `Interpolation`).

### Quantification of the drug interaction type and bottleneck dependency

#### Loewe interaction score

To quantify the drug interaction between a pair of antibiotics, we defined the Loewe interaction score as

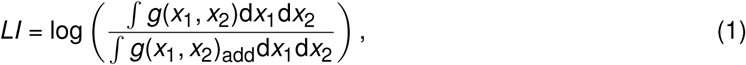

where *g*(*x*_1_, *x*_2_) and *g*_add_(*x*_1_, *x*_2_) are the measured and the predicted additive dose-response surfaces over a 2D concentration field (*x*_1_, *x*_2_), respectively. The score *LI* is a log-transformed ratio of volumes underneath the dose-response surfaces. It is positive for antagonistic and suppressive interactions, 0 for perfectly additive, and negative for synergistic interactions. To avoid imposing arbitrary bounds for classifying a measured interaction as synergistic or antagonistic/suppressive (rather than additive), we performed smooth bootstrapping on a set of ideal additive response surfaces to establish a distribution of interaction indices expected for perfectly additive but noisy surfaces. To achieve this, we generated additive dose-response surfaces for drugs with Hill steepness parameter *n* between 1.8 and 6.6 (obtained as 10% and 90% percentiles of the steepnesses distribution for measured dose-response curves). We estimated the variabilities of measurements *σ*_*v*_ from data from eight replicated dose-response curves with seven replicates per data point and fitting errors *σ*_*f*_ from the slope of all growth rate fits. Both error and variability distributions were well described by log-normal distributions. For each point on the generated surfaces, we added Gaussian noise with standard deviation given 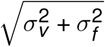, where both *σ*_*v*_ and *σ*_*f*_ were drawn from respective log-normal distributions. We calculated *LI* score for 2000 response surfaces and obtained the distribution shown in Fig. S1D. We determined boundaries separating synergistic and antagonistic *LI* scores (*b*_lower_ and *b*_upper_, respectively) from additive interval as Bonferroni-corrected percentiles (for 5%/28 ≈ 0.18% and 99.82%) of the bootstrapped distribution Fig. S1D). Mean *LI* scores for measured response surfaces falling outside of the interval with these boundaries were classified as synergistic or antagonistic; otherwise, the interaction was classified as additive.

#### Bottleneck dependency score

Similar to *LI*, the bottleneck dependency score *BD* is an integrative quantity that concisely reports on the response-averaged deviation from independence. To calculate this score, the antibiotic and inducer concentrations are first converted into corresponding responses using the induction- and antibiotic dose-response curves (Fig. S3). Mathematically, this means that *r*_*x*_ = *y* (*c*) and *r*_*y*_ = *g*(*in*) for antibiotic and inducer, respectively. In response space, the null-expectation is independence, *i.e.* the expected response is a product of individual responses. Thus, we define the *BD* score as

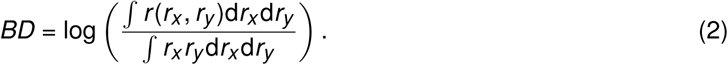

This score is zero when the two perturbations (bottleneck and antibiotic) are independent; it is positive or negative for alleviation and aggravation, respectively. As for the *LI* score, we evaluated the independence interval of *BD* scores by bootstrapping the *BD* score for independent surfaces at given induction and antibiotic dose-response curves. Evaluating the percentiles of such null-distributions gave the boundaries for evaluation of the type of deviation from independence (alleviation or aggravation).

### Growth law-based biophysical model

#### Single antibiotic

The mathematical model used for predicting bacterial growth in the presence of antibiotic combination is an extension of the model presented in Ref. [Greulich *et al.*, 2015]. In-depth analysis of the model will be presented elsewhere. In brief, the model captures the crucial processes of antibiotic binding and transport as well as physiological constraints. We briefly summarize the results for a single antibiotic and its main ingredients. The growth laws are given as

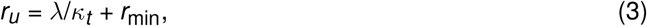

and

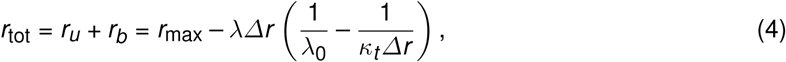

where *r*_*u*_, *r*_*b*_ and *r*_tot_ are concentrations of unbound, bound and total ribosomes. The constants *κ*_*t*_ = *µ*M^−1^h^−1^, *r*_min_ = 19.3 *µ*M, *r*_max_ = 65.8 *µ*M and *Δr* = *r*_max_ – *r*_min_ = 46.5 *µ*M were experimentally determined in Refs. [Scott *et al.*, 2010; Greulich *et al.*, 2015]. Transport of antibiotic is captured by the average flux as *J*(*a*_ex_, *a*) = *p*_in_*a*_ex_ – *p*_out_*a*, where *p*_in_ and *p*_out_ are influx and efflux rates, respectively, and *a* and *a*_ex_ are the intracellular and external antibiotic concentration, respectively. The kinetics of binding of the antibiotic to the ribosome is given as *f* (*r*_*u*_, *r*_*b*_, *a*) = –*k*_on_*a*(*r*_*u*_ – *r*_min_) + *k*_off_*r*_*b*_, where *k*_on_ and *k*_off_ are binding and unbinding rates, respectively, and *K*_*D*_ = *k*_off_/*k*_on_. The fraction of inactive ribosomes *r*_min_ is assumed not to bind antibiotics [Greulich *et al.*, 2015]. The following system of ordinary differential equations (ODEs) describes the kinetics of the system

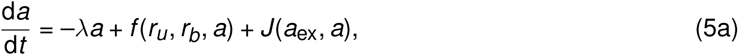

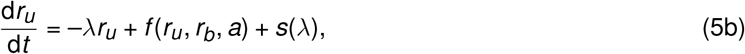

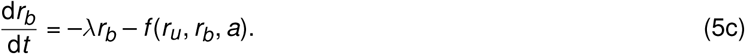

In Eqs. (5) the terms -*λX* (with *X* = *a, r*_*b*_, *r*_tot_) describe effective dilution due to growth and *s*(*λ*) = *λr*_tot_ is the ribosome synthesis rate. In balanced exponential growth all time derivatives in Eqs. (5) vanish and the steady-state solution reads

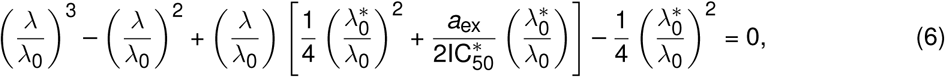

where 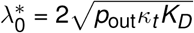 and 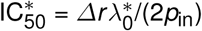. This equation can be recast into

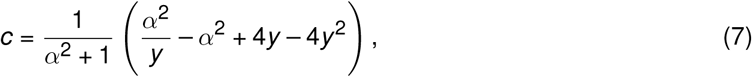

where

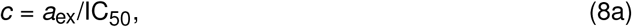

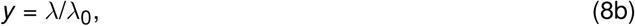

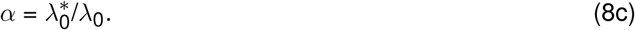

Here, IC_50_ is the concentration required to halve the growth rate (compared to zero drug) and we took into account that 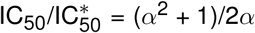. Importantly, the dependence of the relative growth rate *y* on the relative concentration *c* dramatically changes when 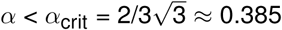, as Eq. (7) exhibits a concentration interval in which growth rate has two stable solutions.

#### Pair of antibiotics

When a pair of antibiotics is considered, additional ODEs are added to describe the binding of individual antibiotics to ribosomes (first binding step) as well as the simultaneous binding of two antibiotics to the already bound ribosome (second binding step):

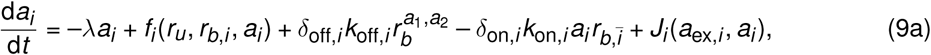

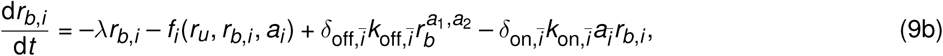

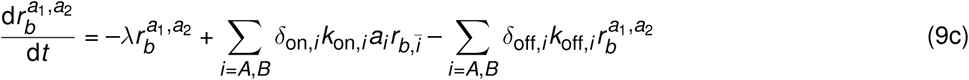

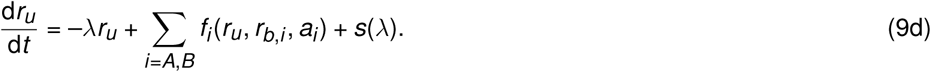

In the system of Eqs. (9), the kinetic parameters and the transport flux and binding functions depend on the antibiotic (indices *i*). The additional terms 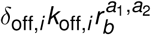 and 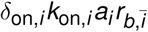 describe the rates of detachment of the *i*-th antibiotic from double-bound ribosomes 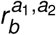 and binding of the *i*-th antibiotic to the ribosome already bound by the other antibiotic *ī*, respectively. The parameter *δ*_*j,i*_ determines the relative changes in rate constants when the other antibiotic is bound. Here, we investigated two cases: independent binding of the two antibiotics, *i.e., δ*_*j,i*_ = 1 and competition *δ*_*j,i*_ = 0, where binding of either antibiotic excludes binding of the other one. The effects of different values of *δ*_*j,i*_ will be presented elsewhere.

We obtained the steady state solution of Eqs. (9) numerically by forward time integration (Mathematica function `NDSolve`). While forward integration requires explicit values of kinetic constants *k*_on_ and *K*_*D*_, the steady state solutions are largely independent of the exact parameter values as long as the parameter ratios *α* and 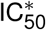 are the same and *k*_on_ ≫ *κ*_*t*_. Upon fitting *α* to the normalized dose-response curves, we fixed *k*_on_ = 100 *µ*M^−1^h^−1^ (which gave consistent results for all dose-response curves). For each dose-response curve, we determined the optimized value of *K*_*D*_ – this was required due to explicit need of parameters in forward integration (Fig. S2). By constraining these parameters, we can calculate the steady state solutions of Eqs. (9).

### Clustering of bottleneck-dependency vectors

We performed clustering of BD vectors projected on a space of lower dimensionality. For dimensionality reduction, we used Principal Component Analysis (PCA). We used the first three principal components which explained *η*_*r*_ ≈ 95.38% of variance. In this projected three-dimensional space, we performed unsupervised agglomerative clustering (Mathematica function `FindClusters`) with cosine distance as a measure of cluster cohesion.

We estimated the p-value of the observed clustering by bootstrapping. We used the Rand index (*RI*) [Rand, 1971] as a criterion for evaluating the difference between clustering results. For example, if *w* is the clustering obtained for the reshuffled sample and clustering *w*′ is obtained for PCA projection of median bottleneck dependency vectors (shown in Figs. 3,S3), then the Rand index is

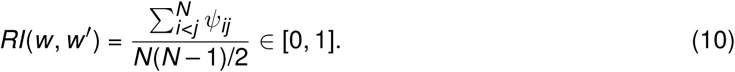

Here, *ψ*_*ij*_ is 1 if the *i*-th and *j*-th data points are either inside or outside of the same cluster and zero otherwise; the denominator is the total number of unique pairs between *N* elements. We generated 10^4^ reshuffled datasets, evaluated *RI* for each dataset and calculated the cumulative distribution function. We evaluated an empirical p-value as

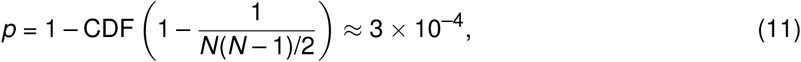

which is an estimate of the probability for obtaining the observed clustering of median BD vectors by chance. The cluster areas shown in Fig. 3 were obtained by smooth bootstrapping of median BD vectors for a given noise statistics, PCA projection and subsequent calculation of the minimal convex hull (Mathematica function `ConvexHullMesh`). The additional response vectors for LAM, TMP and NIT were PCA projected (using Mathematica function `DimensionReduction` obtained for the median values of BD vectors). Note, that the plots in Fig. 3E show projections onto PC1,2 but clustering was performed on first three principal components (Fig. S3).

### Remapping

Our remapping procedure converts inducer concentrations *in* into the concentrations *c* of an idealized antibiotic that precisely targets the translation step controlled by the titrated factor. This requires an induction curve and a dose-response curve: The former is described by an increasing Hill function *g*(*in*), and the latter by solving Eq. (7) for *y*. The conversion between concentrations is formally described as *c* = *y* ^−1^ (*g*(*in*)) at a given *α*, which can be arbitrarily chosen for the idealized antibiotic. When *α* < *α*_crit_, the dose-response curve is bistable and has a region in which more than one response will yield the same concentration – in these cases we consider only the concentration corresponding to the highest stable growth rate as the other solutions are either unstable or will be outcompeted. Further, higher inducer concentrations are remapped to lower antibiotic concentrations and an infinite inducer concentration corresponds zero antibiotic concentration. As this is impractical, we considered all mimicked concentrations (normalized with respect to IC_50_) that are below 0.1 as equivalent to 0.

#### Regularization of surfaces

Strains containing the factor titration platform have mostly very similar antibiotic dose-response curves as the wild-type at maximal inducer concentrations. However, to correct for small deviations, we rescaled the antibiotic concentrations on the antibiotic-inducer grid. The shape of this transformation is derived from equating the responses of two Hill functions with different steepnesses. Consider two Hill functions with Hill exponents *n*_WT_ and *n*_*t*_ for WT and factor-titrating strain, respectively. Then, by equating the responses captured by these Hill functions, we calculated the rescaled relative (with respect to IC_50_) antibiotic concentrations as 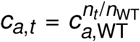. We refer to this conversion as the “power-law transform”. Such regularized surface was then used in remapping.

#### Remapping-based equivalence

Factor deprivation is equivalent to the action of a specific antibiotic if both perturbations can substitute for each other. Upon remapping the inducer concentration, the response surface for an equivalent inducer-antibiotic pair is transformed into an additive response surface. To determine if the deprivation of a specific factor is equivalent to the action of a specific antibiotic, we performed the remapping in tandem with bootstrapping. Bootstrapping assesses the effects of uncertainties in the remapping parameter *α* (obtained from a fit to a drug dose-response curve), artifacts of the response surface over inducer-antibiotic grid and sampling, and inherent noisiness of growth rate determination. We first restricted the dataset to data points with relative growth equal to 0 or above 0.1 with growth rate coefficient of determination *R*^2^ > 0.8. In each round of bootstrapping, the following steps are carried out:

- drawing of a remapping parameter *α* from a normal distribution, centered at the best-fit-value and with standard deviation estimated from fitting, and remapping,
- drawing of a random sample from remapped data points that is of random size (between 75% and 100% of the data set),
- addition of Gaussian noise to the growth rates (estimated from the growth rate fit),
- calculation of the ideal additive surface at a given *α* for comparison, and
- calculation of *LI* score.

This procedure was repeated 100 times for each bottleneck-antibiotic pair and yielded a set of distributions. Each *LI* distribution was then statistically evaluated for being inside the additive interval. We obtained the cumulative distribution function (CDF) for each distribution and we calculated its value on both ends of additive interval (Fig. S1). If either 1 – CDF (*b*_lower_) or CDF (*b*_upper_) is below *p* = 0.05, the pair is considered inequivalent – this is the case in which the remapped surface is unlikely to be additive. For each antibiotic, more than one of the bottlenecks could be statistically equivalent – we thus deemed the bottleneck-antibiotic pair with the highest correlation between average remapped and ideal additive growth rates to be the primary candidate for equivalence of perturbations.

### Quantitative comparison of predicted and measured response surfaces

Both measured and predicted surfaces match along the individual concentration axes, as those were obtained from the fits of dose-response curves. Thus, points corresponding to such measurements are always a good match and in turn increase Pearson correlation invariantly of a potential mismatch in surface segments further away from individual axes. We thus sought an applicable metric that would identify systematic deviations from predicted isoboles.

We developed an “isobole sliding” method in which we determine a mean deviation of points close to some predicted growth rate from measured values. It provides a concise quantitative description of differences between predicted and measured isoboles and identifies the most discrepant areas of the surfaces. For that we systematically move along the (ordered) predicted growth values *g*_*i*_ and select *S* = 20 consecutive points and average their deviations from measured values of growth rate *h*_*i*_. This yields a deviation trajectory *t*(*ĝ*) of a mean deviation as a function of average predicted growth rate

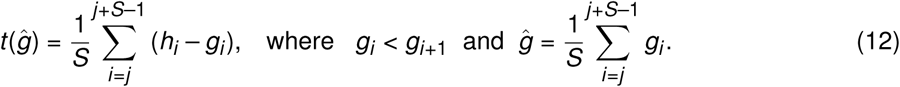

Keeping the number of points *S* in the window fixed allows the comparison between different subsets of the data.

To assess the probability of observing such deviation by chance, we created a benchmark dataset by replacing all measured values with predicted ones to which we added Gaussian noise (estimated from bootstrapped dispersion, but of at least 0.05 relative growth units). For each bootstrapped realization (obtained either by remapping or the biophysical model), we randomly drew a subset of random size (between 75% and 100% of the data set) to estimate the robustness of the prediction with respect to a low number of outliers. We collapsed each isobole sliding trajectory into a single number (*s*) by calculating an maximal deviation, 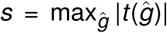, thus yielding a distribution of *s* values for both measured and benchmark trajectory maxima.

Ideally, the distribution of maximal average deviations should either overlap or be below the benchmark distribution. To assess the statistical deviation, we evaluated the CDF of predicted-measured distribution at the 95-percentile of the benchmark distribution. If the value was below 0.05, we rejected the prediction. This method requires that there are no systematic deviations over the whole surface, thus yielding a very stringent criterion for considering a match between two surfaces. Thus, even if two surfaces match qualitatively, isobole sliding might still return a statistically significant mismatch.

To estimate the upper bound of prediction-measurement consistency, we checked for consistency of the measured replicates. For this we considered one of the replicates as a prediction of the other. Doing so, we observed that twenty-one out of twenty-eight (75%) surfaces act as statistically significant predictions for one another. This serves as an approximate upper bound for how many predictions-measured pairs can be at most expected to match at the given experimental variability.

#### Assessment of predictive power

At this point we can assess the consistency of predictions. Using the method described above, we evaluated both independent and competitive binding schemes for their congruence with measured surfaces. The scheme that led to the distribution with the smallest mean maximal deviation, was considered as best-match. However, both schemes can yield a good match – by asking how many of the schemes yield a match in both replicates, we obtain an estimate for a fraction of correct predictions (Fig. S2). By counting in how many cases at least one of the schemes yields a match between replicates, we find that sixteen out of twenty-eight interactions can be accounted for by a biophysical model.

Applying isobole sliding to the prediction of remapping shows that even small quantitative deviations will lead to discarding of the prediction (Fig. S5). However, counting additionally explained interactions by remapping (TET-CRY, TET-FUS, KSG-CHL, CRY-KSG) increases the total tally of explained interactions to twenty out of twenty-eight (≈71.4%), which is below the estimated self-consistency bound of 75%. As discussed above, qualitative matches are not included in this metric.

### TASEP model of translation within growth law framework

There are several specific differences between the classical open TASEP system and translation in the context of the bacterial cell. Firstly, the pool of ribosomes is finite and variable in size (as dictated by the growth laws). Secondly, the ribosomes span over more than one site – it occupies *L* ≈ 16 codons [Kang and Cantor, 1985]. Thirdly, steps in translation are mediated by translation factors that bind to the ribosome in a specific state and (stochastically) push the ribosome into another state. The rates depend on the abundance of ribosomes in a specific state and the abundance of the factor catalyzing the step. Thus, the rates, which are kept fixed in the classical TASEP, become variable and system-state dependent.

#### Mathematical framework

##### Analytical results for TASEP of extended particles

In the absence of ribosome pausing, established analytical results for the TASEP of extended particles can be used [Klumpp and Hwa, 2008; Lakatos and Chou, 2003; Shaw *et al.*, 2003; Zia *et al.*, 2011]. If the release of ribosomes at the end of the transcript is not limiting, two different regimes of ribosome traffic exist, namely the initiation- and translocation-limited regime. These regimes are separated by a non-equilibrium phase transition. The current of ribosomes *J* in the two regimes is given by:

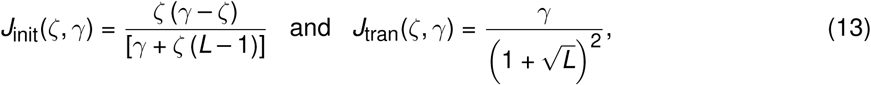

where *ζ* and *γ* are initiation and translocation attempt-rates, respectively. The ribosome (coverage) density *ρ* reads:

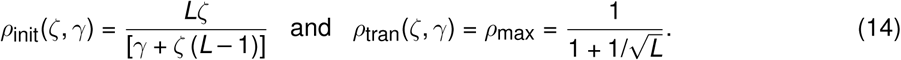

The elongation velocity *u* depends both on the current and the ribosome density *ρ*_*r*_ = *ρ*/*L* via *u* = *Js*/*ρ*_*r*_, where *s* is the step size (1 aa). This in turn yields

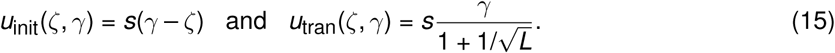

##### Distribution of ribosomes across different classes

The total ribosome concentration *r*_tot_ is

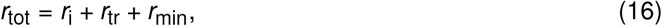

where *r*_i_ and *r*_tr_ are the concentrations of non-initiated and translating ribosomes, respectively. Translating ribosomes are distributed across numerous mRNA transcripts in the cell and their concentration can be written as:

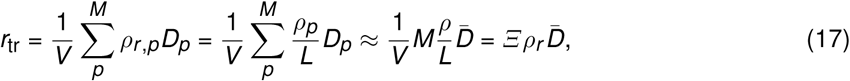

where *D*_*p*_ and *ρ*_*r,p*_ are the length and ribosome density of the *p*–th transcript, respectively, *M* is the total number of transcripts and *V* the cell volume (*Ξ* = *M*/*V* is the concentration of transcripts). The density of ribosomes *ρ*_*r*_ = *ρ*/*L* is a TASEP-derived quantity and depends on the initiation attempt rate *α* and translocation attempt rate *γ*. In the last step, we assumed for simplicity that the density of ribosomes across the transcripts does not vary significantly between transcripts. However, if transcripts do differ in their ribosomes densities, the ones with higher densities will enter the translocation limiting regime (in which traffic jams form) already at a smaller decrease in translocation attempt rate. If those transcripts code for essential genes, this will correspondingly lead to a decrease in growth rate already at such smaller decreases in translocation attempt rate. Such traffic jams would still be relieved by lowering initiation rate even though traffic jams have not developed on all other transcripts. Thus, the qualitative conclusions of the analysis below would still hold, but the results would be quantitatively different. However, taking differences between transcripts into account would require explicit modeling of individual transcripts and is beyond the scope of this work. Assuming similar ribosomes densities allows replacement of the sum with 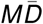, where 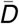 is the average length of transcripts being translated; the proteome-weighted average length is 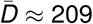 [Milo and Phillips, 2016].

The growth rate is proportional to the elongation velocity of ribosomes along the transcript *u*(*α, γ*) and to the number of translating ribosomes. However, there is a limit for the maximal elongation rate *u*_max_ because other processes (*e.g.*, charged tRNA delivery) become limiting at some point in a given nutrient environment. We estimated the maximal elongation rate from the Michaelis-Menten-like relation between RNA/protein (*R*/*P*) and translation rate obtained in Ref. [Dai *et al.*, 2016]: *u* = *k*_el_(*R*/*P*)/[(*R*/*P*) + *K*_el_], where *k*_el_ = 22 aa/s and *K*_el_ = 0.11. We calculated the theoretical 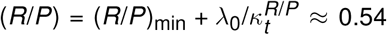, where 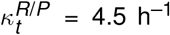 and (*R/P*)_min_ = 0.09[Scott *et al.*, 2010]. Plugging this (*R*/*P*) into the Michaelis-Menten function for the translation rate, we obtain *u*_max_ ≈ 18 aa/s. Thus, the growth rate is given as

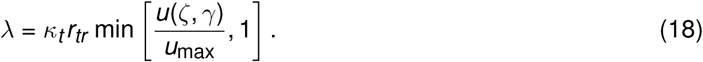

However, the growth rate feeds back into the total ribosome concentration via the growth law as

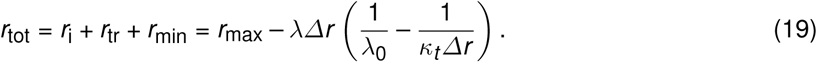

We can estimate *Ξ* at *λ*_0_ as

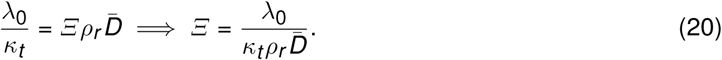

##### Factor-dependent translocation attempt rate

The ribosome will perform a specific step only when the associated factor is bound to it: the step-attempt rate is proportional to the probability *P*_*b*_ of the ribosome being bound by a factor. This probability can be calculated by assuming a population of elongation factors with concentration *c*_ef_ = *c*_ef,b_ + *c*_ef,n_ and translating ribosomes *r*_tr_ = *r*_tr,b_ + *r*_tr,n_, where the indices *b* and *n* denote the factor-bound and unbound subpopulations, respectively. Binding is described by

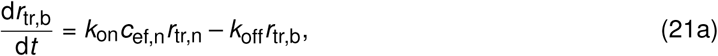

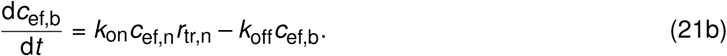

Solving for steady state, noting that *r*_tr,b_ = *c*_ef,b_ and defining *K*_*D*_ = *k*_off_/*k*_on_ we obtain the probability for a ribosome to be bound as

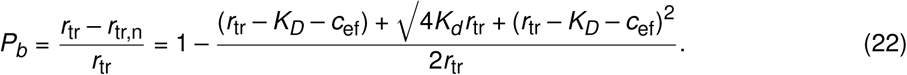

The binding constant of EF-G to the ribosome complex I (pre-translocation analog with N-Ac-dipeptidyl-tRNA at the A-site and deacylated-tRNA in the P-site) [Yu *et al.*, 2009] is *K*_*D*_ = 0.27 ± 0.02 *µ*M; we used this value in our calculations. In the case of WT regulation there are ∼ 0.83 EF-G molecules per ribosome and the expression of the factor is coupled to the ribosome number (*i.e.*, their ratio is constant) [Dai *et al.*, 2016].

##### Factor-dependent initiation attempt rate

Successful initiation events are not limited to a single *L*-codon long slot on a mRNA (that can be free or occupied) but can occur on any transcript; and only the factor-bound ribosomes can attempt an initiation event. Thus, the initiation rate can be described by Michaelis-Menten kinetics:

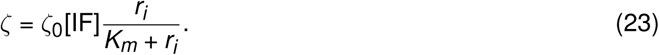

We can estimate *K*_*m*_ from kinetic rates determined by Milon *et al* [Milon *et al.*, 2012] where the free 30S subunit is bound (almost simultaneously) by IF1 and IF2 with rate (2–10)×10^2^ *µ*M^−1^s^−1^ and dissociates at rate 30 s^−1^. From these values, we estimate *K*_*m*_ ≈ 0.05 *µ*M.

##### Estimation of model parameters

It is useful to estimate if WT translation is in the initiation or translocation limited regime, which we can obtain from the average ribosome density. We can estimate the ribosome density as *ρ*_*r*_ = 3*β*_*r*_ *N*_*r*_ /(*r*_*m*_*t*_*m*_), where *N*_*r*_, *β*_*r*_, *r*_*m*_ and *t*_*m*_ are the number of ribosomes, the fraction of active ribosomes, the rate of mRNA synthesis per cell, and the average mRNA life-time, respectively [Bremer and Dennis, 1996]. The fraction of translating ribosomes *β*_*r*_ is estimated from fitting a Hill function to data from Ref. [Dai *et al.*, 2016] (Fig. S6). For higher growth rates, the relation between growth rate and (calculated) ribosome density linearizes; extrapolating to *λ*_0_ = 2.0 h^−1^, we obtain *ρ*_*r*_ ≈ 0.042 (Fig. S6), which yields *Ξ* ≈ 3.7 *µ*M. For cells grown in LB, the average number of transcripts per cell was measured as *N*_mRNA_ ≈ 7800 [Bartholomäus *et al.*, 2016]. To estimate the mRNA concentration, we use *Ξ* = *N*_mRNA_/*V*_cell_ = (*N*_mRNA_/*m*_dry_) × (*m*_dry_/*m*_wet_) × (*m*_wet_/*V*_cell_), where *m*_dry_/*m*_wet_ ≈ 1/3.1 and *m*_wet_/*V*_cell_ ≈ 1.09 g/mL are growth-rate independent quantities (see SI of Ref. [Greulich *et al.*, 2015]). We obtained the dry mass of the cell at *λ* = 2.0 h^−1^ by extrapolating from measured data at various growth rates [Bremer and Dennis, 1996] as *m*_dry_ ≈ 1.01 pg/cells (Fig. S6) which in turn yields *Ξ* ≈ 4.5 *µ*M. This value differs from the estimate above by ≈ 22%.

The estimated ribosome density is *ρ*_*r*_ ≈ 0.042, which is lower than the maximal attainable ribosome density of 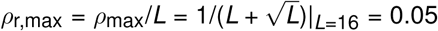. Thus, translation in the WT is likely in the initiation-limited regime. Thus, the equations for ribosomal density and elongation velocity for the initiation limiting regime are used to estimate the apparent initiation and translocation attempt rates:

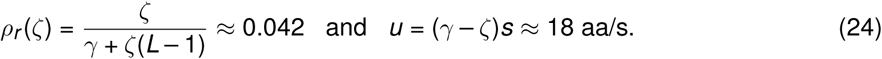

The apparent rates are *γ* ≈ 20.3 s^−1^ and *ζ* ≈ 2.3 s^−1^. This allows us to estimate *γ*_0_ = *γ*/*P*_*b*_, where we note that *c*_ef,WT_ ≈ 43.0 *µ*M (estimated from 0.83 × 51.9 *µ*M where the ribosome concentration is calculated from the growth law). Next, we estimate the number of translating ribosomes from Eq. (18) as 32.6 *µ*M, which yields *P*_*b*_ ≈ 0.98 and finally *γ*_0_ ≈ 20.7 s^−1^. We further note that there are 0.3 IF2 molecules per ribosome [Bremer and Dennis, 1996], implying [IF]_WT_ ≈ 15.6 *µ*M, from which we estimate *ζ*_0_ ≈ *ζ*/[IF]_WT_ ≈ 0.15 *µ*M^−1^s^−1^.

With these parameter values, our model is fully defined and the growth rate is calculated (Mathematica function `NSolve`) as its output based on the concentration of translation factors. To verify the impact of unperturbed ribosome density *ρ*_*r*_ (one that supports maximal growth rate at saturating factor concentrations), we systematically calculated the response surfaces for different values of *ρ*_*r*_ between 0.001 and 0.049 (Fig. S6). With decreasing unperturbed *ρ*_*r*_, the concentration of mRNA *Ξ* increases according to Eq. (20). When 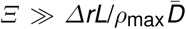, the traffic jams of ribosome are not possible anymore as there are too many mRNAs that can carry more ribosomes than available. The critical unperturbed ribosome density is *ρ*_r,crit_ = *λ*_0_/(*κ*_*t*_ *Δr*) × *ρ*_r,max_ (Fig. S6).

#### Effect of mRNA growth-rate dependence

The concentration of mRNA could in principle be growth rate-dependent. However, direct dependence of mRNA as a function of the growth rate is difficult to estimate from existing literature as total RNA is mostly composed of rRNA and tRNA [Dai *et al.*, 2016; Scott *et al.*, 2010]; estimation of the mRNA fraction is thus prone to errors. However, if we assume proportionality between ribosome and mRNA concentration, a simplified form can be written down as *Ξ* = *Ξ*_0_*r*_tot_/*r*_tot,0_, where *Ξ*_0_ and *r*_tot,0_ = *r*_min_+*λ*_0_/*κ*_*t*_ are the estimates of mRNA concentration from the previous section and total ribosome concentration in the unperturbed case, respectively. Plugging this dependence into the model does not qualitatively change the suppressive interaction between inhibition of initiation and translocation (Fig. S6). In this scenario, the increasing number of mRNA transcripts partially alleviates the densification of ribosomes on transcripts. However, the over-all increasing number of translating ribosomes sequesters the elongation factors – this effect is still alleviated by lowering the initiation rate and in turn the density of ribosomes.

#### Rescue mechanisms and inefficiency of direct response to translocation inhibition

Bacteria have evolved rescue mechanisms for stalled ribosomes (tmRNA, ArfA and ArfB). However, these mechanisms are mostly aimed at the rescue of ribosomes that were stalled due to limiting supply of building blocks or those in non-stop complexes. Former is an unlikely scenario during translocation limitation: as the building blocks are under-consumed, non-stop complexes can form via the formation of damaged or truncated mRNA (*e.g.*, via cleavage by RNases) or via collision-induced frameshifts [Simms *et al.*, 2019; Keiler, 2015]. However, the tmRNA pathway requires an empty A-site on the ribosome, which is occupied in the pre-translocation complex, thus hindering the rescue initiation. Likewise, the ArfA pathway is hindered by an occupied A-site – it requires release factor 2 to bind to the A-site of the ribosome to initiate premature release and recycling. ArfB on the other hand, can recover the lack of tmRNA and ArfA pathways only when heavily overexpressed [Chadani *et al.*, 2011] and is considered ineffective in the WT regime. In sum, established rescue mechanisms are unlikely to recover stuck ribosomes and we therefore omit these mechanisms from the analysis.

Additionally, the cell could have an initiation-inhibiting mechanism in place as a response to translocation inhibition. However, the observed responses of bacteria to translation inhibition show global derepression of the translation machinery by reducing the levels of ppGpp. Besides the upregulation of all translation components mentioned in the main text [Maaløe, 1979; Gordon, 1970; Blumenthal *et al.*, 1976; Furano and Wittel, 1975], an additional effect of lower levels of ppGpp is a direct increase of initiation. The catalytic function of the initiation factor is lowered when ppGpp levels are high, and higher when ppGpp is reduced [Milon *et al.*, 2006]. These arguments show that an alleviating response of translocation inhibition by either rescue mechanisms or by direct down-regulation of initiation is unlikely.

## 8 Supplementary Information

**Table S1: Chemicals used in this study.** Table contains chemical names and purpose categories, catalog codes and vendor information.

**Table S2: Oligonucleotides used in this study.** Spreadsheet contains primer names, sequences, templates and brief description of use. Spreadsheet is divided into tabs, each corresponding to the aim of a specific cloning step.

**Figure S1:**
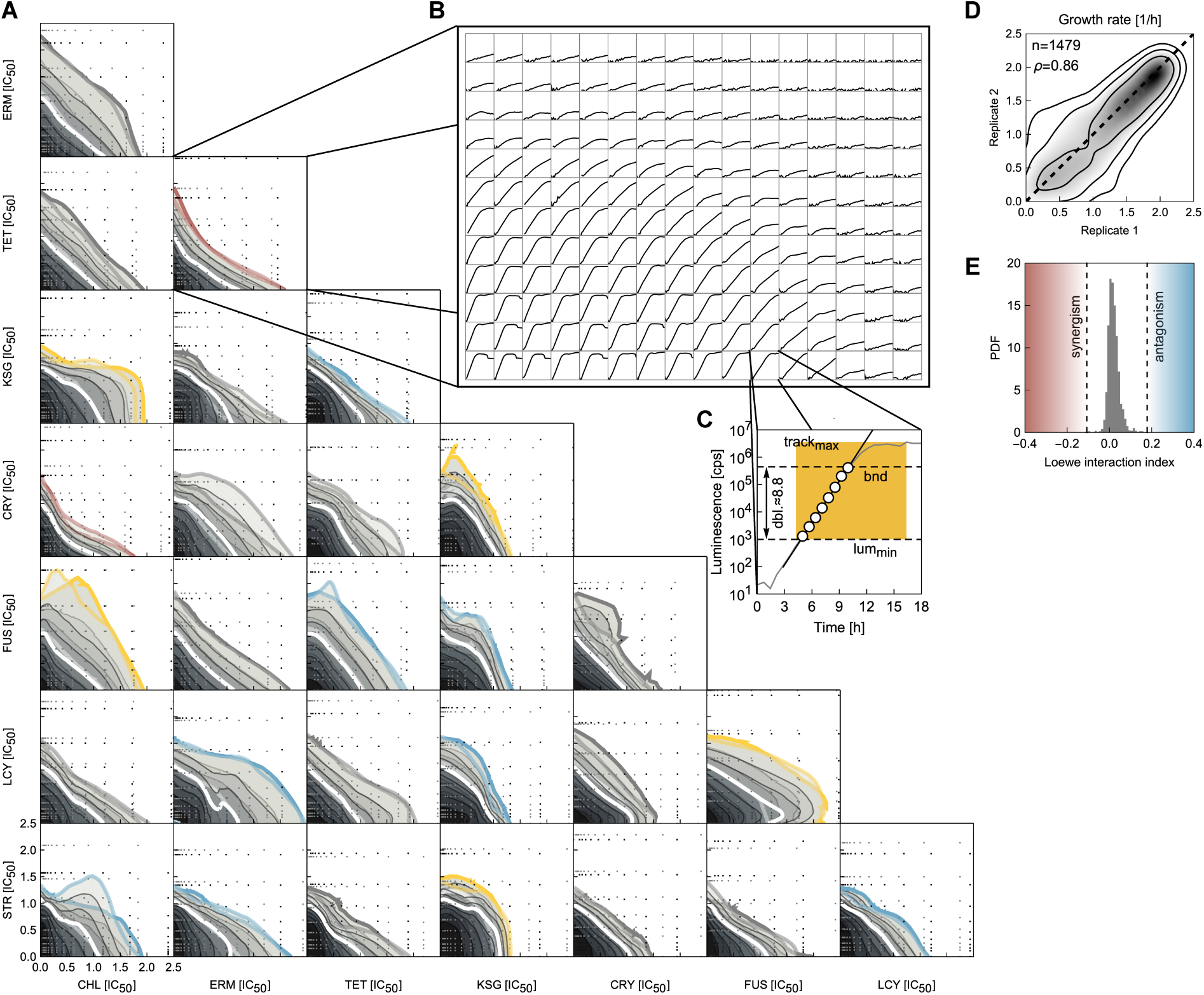
All dose-response surfaces and examples of growth curves. **(A)** Duplicates of dose-response surfaces for all 28 antibiotic pairs. Due to small, but systematic variability in concentrations between replicates done on different days, we rescaled concentration axes with respect to the IC_50_. Dose-response surfaces were smoothed using LOESS (Methods). Black and gray dots denote measured points from different experiments. Isoboles from duplicates are in high agreement; small deviations are caused by occasional outliers that skew the isoboles. As the dose-response surface was measured over a 12×16 grid, the duplicates change the drug axes (12×16⟶16×12) on different days to check for effects coming from spreading the measurements over different plates. **(B)** An example of growth curves over a 12×16 grid. Note, that here the concentrations change between wells in a geometric manner, *i.e.* the ratio between concentrations in neighboring wells is fixed. **(C)** Exemplary growth curve and details of the fitting procedure. The growth rate is determined by fitting a line in the regime of exponential growth. The determination of this regime in the growth curve is carried out automatically; procedure: (i) check if the maximum value of luminescence is above the lower bound of the fitting interval lum_min_ = 10^3^ cps and take points before the maximum, (ii) take points that are the latest to rise over lum_min_, (iii) determine the upper limit (bnd) of the fitting interval to be either ten-fold above the lum_min_ (guaranteeing log_2_ 10 ≈ 3.3 doublings of fitting interval) or eight-times less than the track maximum (three doublings away from saturation) and (iv) fit a line to the log-transformed values of the luminescence signal if there are at least three data points. If lum_min_ is not exceeded, the well is counted as having no growth; if any of the other criteria is not fulfilled, growth is characterized as undetermined. **(D)** Reproducibility of absolute growth rate measurements between replicates. The smooth kernel representation of replicate measurements (Mathematica function `SmoothKernelDistribution`), performed on different days and different plate arrangements, demonstrates a good agreement overall. Only non-zero growth rates of sufficient quality (*R*^2^ > 0.5 and relative error < 0.5) are included. **(E)** Distribution of Loewe interaction indices of noisy additive surfaces for pairs of drugs with different steepnesses, as obtained by bootstrapping. Note, that this reveals a slight bias towards antagonism.

**Figure S2:**
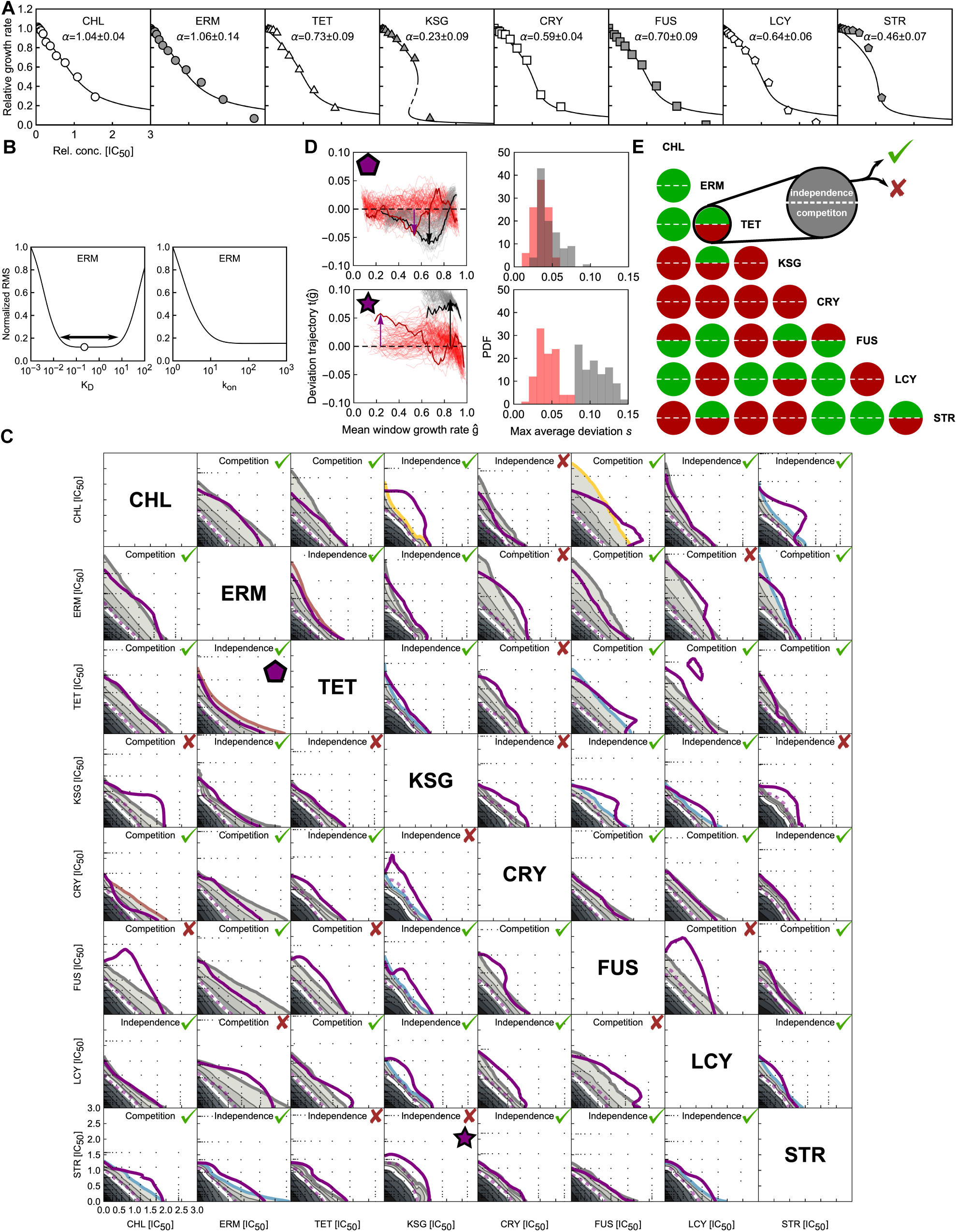
Details of the biophysical model for pairwise antibiotic combinations. **(A)** Average dose-response curves with best fit model for individual antibiotics. Inset denote the corresponding antibiotic and best-fit steepness parameter *α* with standard error. Dose-response curves are predominantly shallow for our selection of antibiotics, *i.e., α* > *α*_crit_. Dashed segment of KSG dose-response curve represents an unstable solution. **(B)** Example of an effect of numerical parameters (*K*_*D*_ and *k*_on_) on root-mean-square error (with respect to the experimental data). Parameters are required for forward time integration. Root-mean-square error was normalized with respect to the maximal error in the scanned interval. Effective dissociation constant *K*_*D*_ exhibits roughly two orders of magnitude wide plateau (double-headed arrow; minimum is denoted by a circle). First order binding rate constant *k*_on_ does not exhibit a plateau but rather flattens out – consistently with the requirement that *k*_on_ ≫ *κ*_*t*_. **(C)** All predictions for replicated measurements. Predicted surface is show in full; overlaid thick and dashed purple isobole denote 20% and 50% isobole, respectively, of the measured surface. Each prediction is evaluated for goodness of prediction as described in Methods. Check-mark and cross denote a match and mismatch, respectively. Inset text denotes the best-matching binding scheme. **(D)** Illustration of isobole sliding method. Left: two examples of deviation trajectories *t* (*ĝ*) for ERM-TET (pentagram) and KSG-STR (five-point star). Thin gray and red lines present hundred bootstrapped repetitions of measured and benchmark trajectories. Two trajectories (thick black and red lines for measured and benchmark, respectively) are highlighted. Black and purple arrows denote maximal deviation of the trajectory from zero for measured and benchmark trajectory. Length of the arrow is max average deviation *s*. Right: all *s* values from bootstrapped repetitions are collected in the histogram. Pair of ERM-TET offers a better match with benchmark distribution compared to KSG-STR. **(E)** Performance of all schemes against the measurements. Upper and bottom half of each circle denote independence or competition, respectively, as denoted. Green and red color denote match and mismatch, respectively.

**Figure S3:**
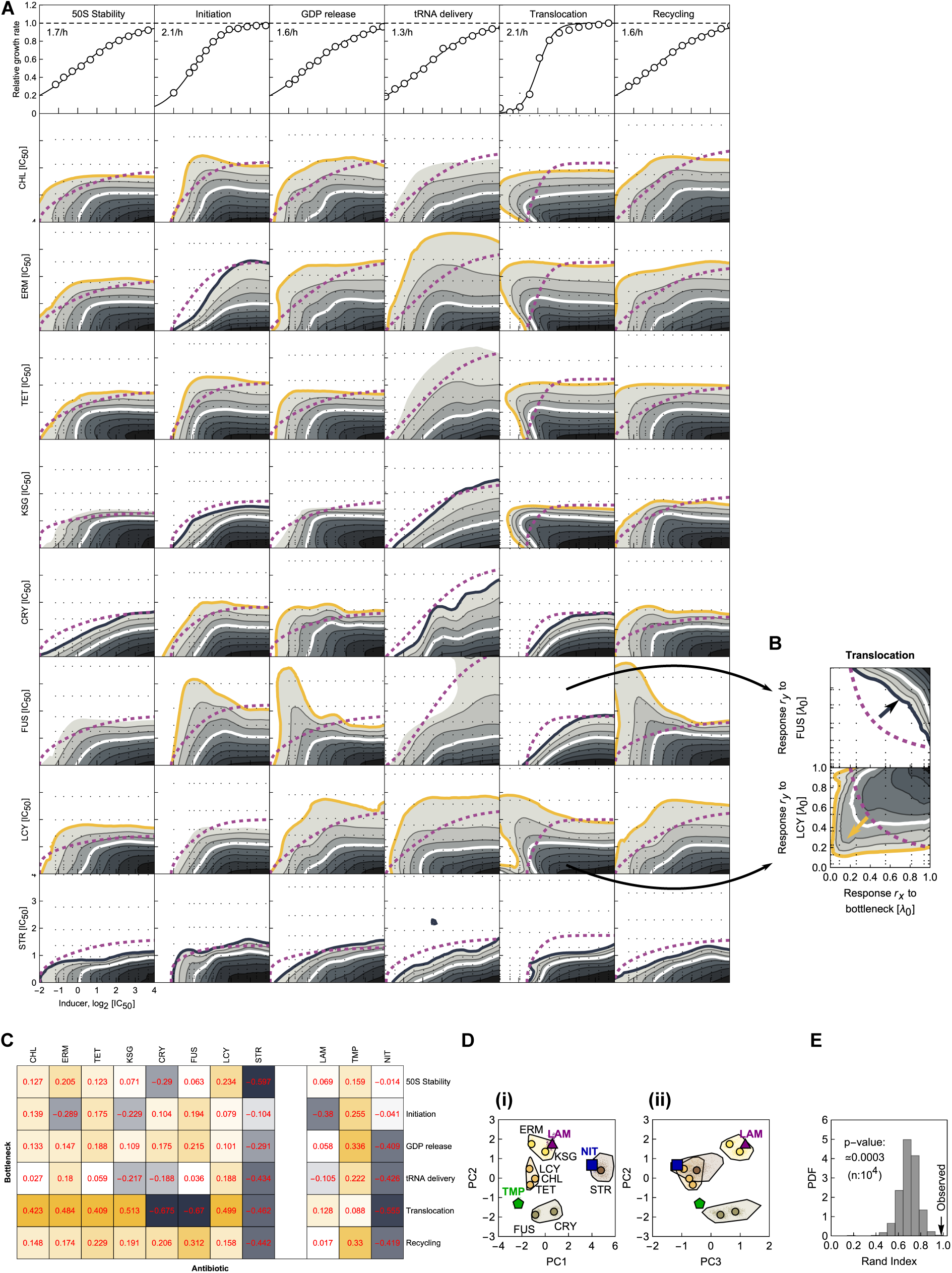
Bottleneck-antibiotic dose-response surfaces and functional classification. **(A)** Dose-response surfaces for all bottleneck-antibiotic pairs. Surfaces were smoothed using LOESS (Methods). Note the different characters of deviations from independence. **(B)** Examples of response surfaces over response-response grid. In the response space (*r*_*x*_, *r*_*y*_), independence is defined as *r*_*x*_ *r*_*y*_. Logarithm of the ratio of volumes underneath the measured and independent surface yields a deviation index. For every antibiotic, six bottleneck dependencies together yield a bottleneck dependency vector. **(C)** Values of bottleneck dependencies for all bottleneck-antibiotic pairs. **(D)** Projection of bottleneck dependencies on PCA vectors. **(i)** As in Fig. 3E. (**ii**) Projection on PCA vectors PC2 and 3. Note the separation of clusters in both projections. **(E)** Bootstrapped clustering of randomized vectors yields a series of clustering results. With these clustering results at hand, we calculate the Rand index *RI*(*w*, *w*′). From the distribution of *RI*(*w*, *w*′), we estimate the empirical cumulative distribution function and corresponding p-value [Eq. (11)] for the clustering result in Fig. 3E.

**Figure S4:**
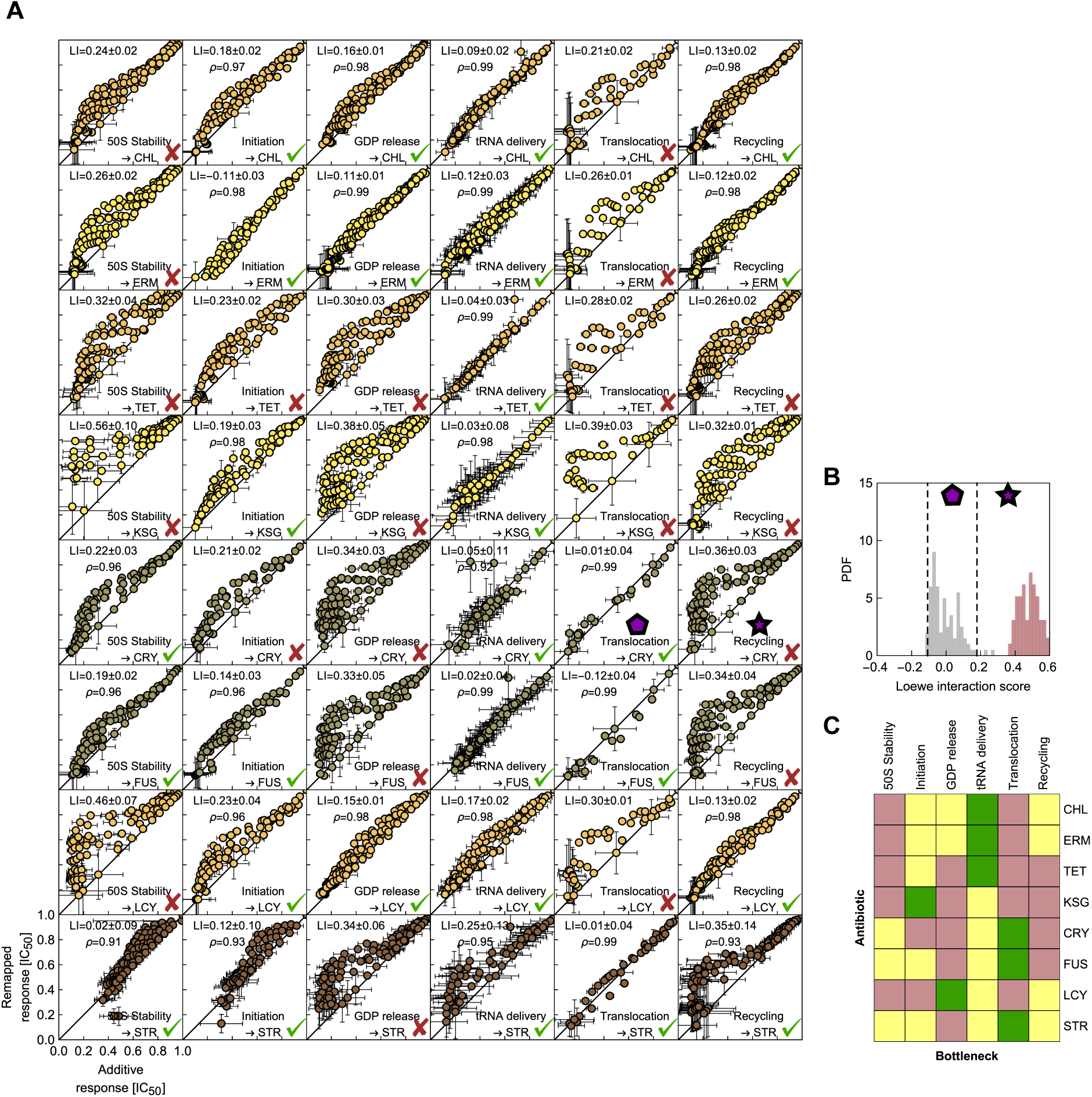
Remapping-based assessment of primary mode of action. **(A)** Scatter plots of growth rates expected for additivity and obtained by self-remapping (Methods). *LI* was statistically compared to the boundaries of the additive interval. Green check marks denote that *LI* did not fall outside of the additivity interval; in these cases, the rounded correlation *ρ* is reported. A good agreement with the additive expectation suggests equivalency of antibiotic and genetic perturbation. **(B)** Examples of histograms of *LI* for CRY in combination with a translocation and recycling bottleneck [see matching pentagon and star in (A)], respectively. **(C)** Color-coded sequential evaluation of equivalence between bottleneck and translation inhibitor. Red and yellow denote that *LI* was outside or inside of the additive interval, respectively. From the cases in which the *LI* is statistically inside the additive interval, the case with highest correlation was chosen as the putative primary mode of action (green). This approach correctly identified the mode of action for all cases in which it is known from literature (CRY, FUS, STR, KSG and TET).

**Figure S5:**
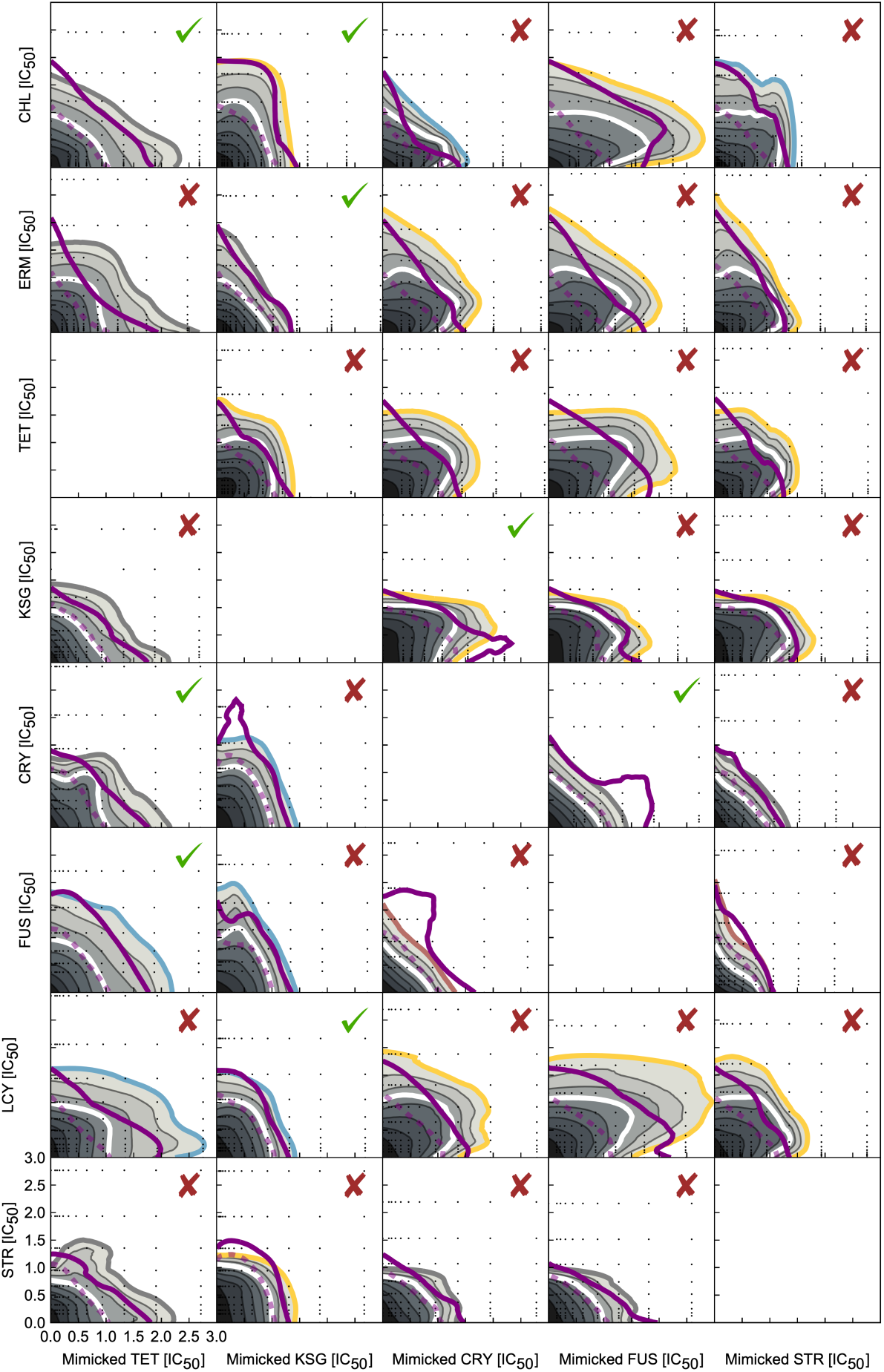
All possible predictions from perturbations of equivalent effects. Predicted surface obtained by remapping is show in full; overlaid thick and dashed purple isobole denote 20% and 50% isobole, respectively, of the measured surface. Each prediction is evaluated for goodness of prediction as described in Methods. Check-mark and cross denote a match and mismatch, respectively.

**Figure S6:**
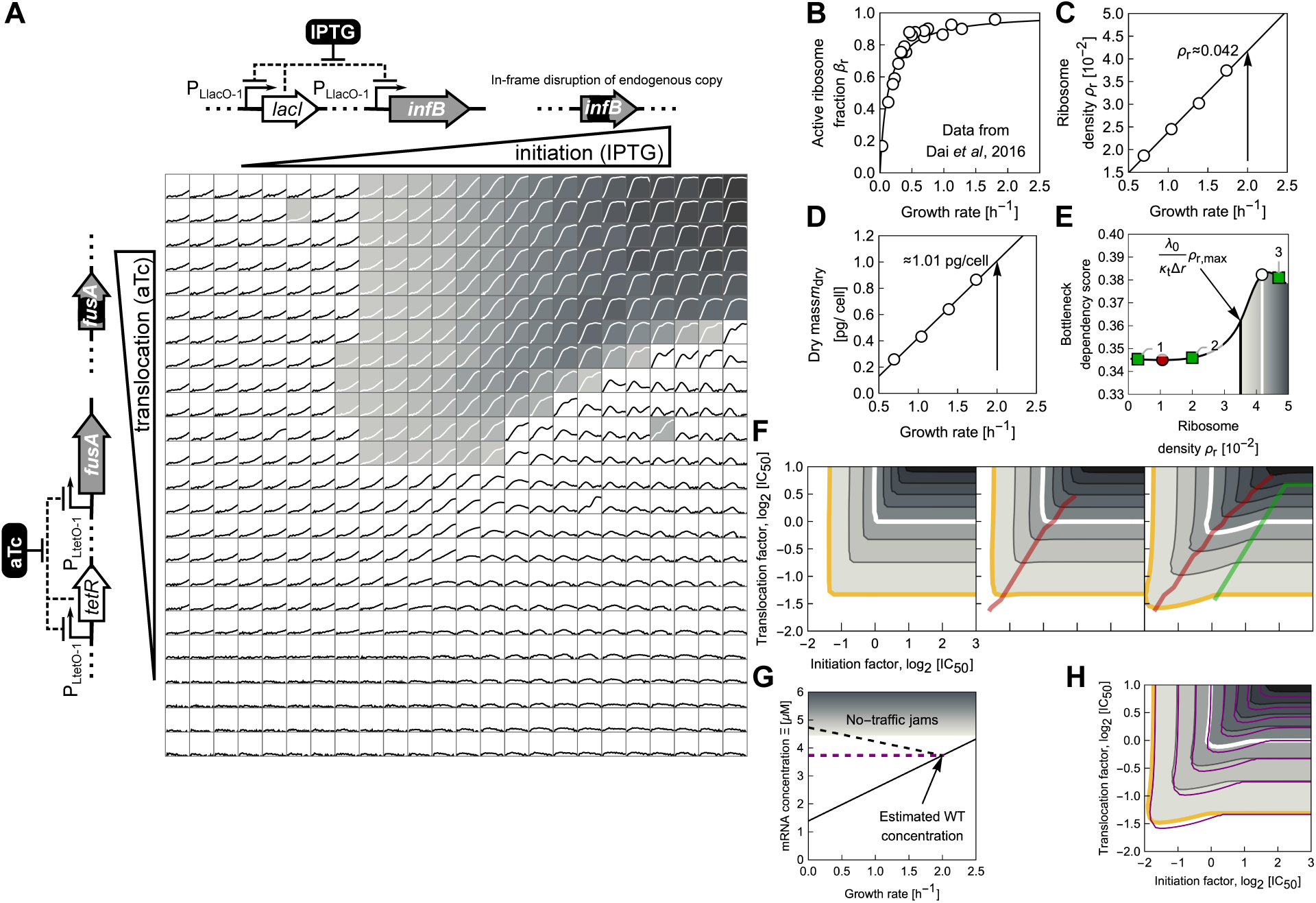
Double titration platform and model analysis. **(A)** Schematics represent the genetic elements of double titration control: negatively auto-repressed transcription factors *lacI* and *tetR* that control the expression of initiation factor *infB* and elongation factor G *fusA*, respectively; expression is dependent on the shown inducers (IPTG and aTc). The grid shows the growth curves for the response surface in Fig. 6. Different shades of gray show the growth rate. Only fits of good quality and with growth rates above 0.199 are included. **(B)** Active ribosome fraction as a function of growth rate in different nutrient environments. Data is from Ref. [Dai *et al.*, 2016]. The solid line represents a best-fit Hill function (*x* /*a*)/ [1 + (*x* /*a*)], where *a* ≈ 0.12 h^−1^. **(C)** Calculated ribosome density *ρ*_*r*_ = 3*β*_*r*_ *N*_*r*_/(*r*_*m*_*t*_*m*_). The arrow denotes the density for *λ*_0_ = 2.0 h^−1^. Solid line shows best fit. **(D)** Dry mass measurements from Ref. [Bremer and Dennis, 1996] and best-fit linear function (solid line). Arrow denotes the density for *λ*_0_ = 2.0 h^−1^. **(E)** Impact of varying the initial *ρ* _*r*_ on resulting bottleneck dependency score. Numbered green squares correspond to the examples showcased in (F). The white circle shows the result for the estimated value of WT *ρr* ≈ 0.042. The red circle shows the point (*ρr* ≈ 0.0106) where first derivative becomes positive and the BD score starts increasing. The solid vertical line shows the critical value *λ*_0_*ρ*_r,max_/(*κ*_*t*_ *Δr*) above which traffic jams due to translocation limitation can form. **(F)** Response surfaces for *ρ*_*r*_ values shown in (E). For *ρr* ≪ 0.01 the ridge line (red; defined by the concentration of initiation factor that supports the highest growth rate at a given concentration of translocation factor) is not well defined, and tends towards high concentrations of initiation factor. For *ρ*_*r*_ > 0.1, the ridge line moves towards the “corner” of the response surface. After the value *λ*_0_*ρ*_r,max_/(*κ*_*t*_ *Δr*) is surpassed, traffic jams develop when the translocation rate is sufficiently low. **(G)** Two models of mRNA concentration dependence. Black lines denote the dependence of mRNA on growth rate if the co-regulation between total RNA and mRNA (Methods) is assumed; solid and dashed lines correspond to variation of the nutrient quality and translation perturbation, respectively. The arrow denotes the estimated mRNA concentration for cells grown in LB (Methods); this concentration is assumed constant (dashed purple line) in the model shown in the main text. If the mRNA concentration exceeds 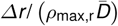, traffic jams do not develop. Elongation factors are still sequestered as the number of translating ribosomes increases, which in turn decreases the growth rate. **(H)** Direct comparison of model predictions. Prediction with growth-dependent mRNA concentration *Ξ* is depicted in full gray-scale tones; isoboles from the prediction assuming a constant pool of mRNA are shown in purple. Both results are qualitatively equivalent.

